# Capsule-Based Single-Cell Genome Sequencing

**DOI:** 10.1101/2025.03.14.643253

**Authors:** Dustin B. Mullaney, Samantha R. Sgrizzi, David Mai, Ian Campbell, Yuqi Huang, Andrius Šinkūnas, D. Lucas Kerr, Valentino E. Browning, Helen E. Eisenach, Jeremiah Sims, Eva K. Nichols, Christopher P. Lapointe, Yasuhiro Amimura, Kelley Harris, Rapolas Žilionis, Sanjay R. Srivatsan

## Abstract

Single-cell genomics methods have unveiled the heterogeneity present in seemingly homogenous populations of cells, however, these techniques require meticulous optimization. How exactly does one handle and manipulate the biological contents from a single cell? Here, we introduce and characterize a novel semi-permeable capsule (SPC), capable of isolating single cells and their contents while facilitating biomolecular exchange based on size-selectivity. These capsules maintain stability under diverse physical and chemical conditions and allow selective diffusion of biomolecules, effectively retaining larger biomolecules including genomic DNA, and cellular complexes, while permitting the exchange of smaller molecules, including primers and enzymes. We demonstrate the utility of SPCs for single cell assays by performing the simultaneous culture of over 500,000 cellular colonies, demonstrating efficient and unbiased nucleic acid amplification, and performing combinatorial indexing-based single-cell whole genome sequencing (sc-WGS). Notably, SPC-based sc-WGS facilitates uniform genome coverage and minimal cross-contamination allowing for the detection of genomic variants with high sensitivity and specificity. Leveraging these properties, we conducted a proof-of-concept lineage tracing experiment using cells harboring the hypermutator polymerase ε allele (POLE P286R). Sequencing of 1000 single cell genomes at low depth facilitated the capture of lineage marks deposited throughout the genome during each cell division and the subsequent reconstruction of cellular genealogies. Capsule-based sc-WGS expands the single-cell genomics toolkit and will facilitate the investigation of somatic variants, resolved to single cells at scale.

## Introduction

Central to biological life is the distinction between self and non-self, inside and outside. This organizational principle manifests across all of life’s scales, from cellular compartments to tissues and whole organisms. At the cellular level, compartmentalization is governed primarily by selectively permeable membranes that actively and passively regulate the flow of materials and information. Passive processes include the diffusion of dissolved gases, hydrophobic molecules, and water through precisely structured protein pores, while active mechanisms transport ions, hydrophilic nutrients, and macromolecules. These transport processes maintain a dynamic equilibrium essential for cellular function. The precise biochemical milieu established by this compartmentalization facilitates not only the concentration of resources and the expulsion of waste, but also organizes the complex biochemical reactions that underpin life.

Experimentally dissecting complex biological systems necessitates methods that harness and preserve this intrinsic compartmentalization. Specifically, measuring the concurrence of and dependency between entities located within the same cell provides insight into the logic underlying biological systems (*1*). One effective strategy involves recreating perturbations of cellular components *in situ* (*2–4*), enabling a view into the roles that individual genes play in biological processes. Such methods have greatly benefited from multiplexing, exemplified by single-cell DNA sequencing technologies, which transform individual cells into isolated compartments, each simultaneously serving as both the experimental container and measurement unit (*5*).

However, traditional compartmentalization methods have limitations. Microtiter wells, despite their widespread use, struggle to scale due to the challenge of precisely manipulating small volumes and actuating over small physical distances (*6*). Droplet microfluidics offers scalability by creating isolated droplets stabilized by emulsifiers (*7*, *8*); yet, these droplets are static, non-dynamic compartments incapable of emulating the dynamic exchange characteristic of living cells. Hydrogel capsules comprising a shell and core begin to resemble the ideal experimental container. Capsules are thin walled, hollow structures formed with droplet microfluidics that enable the equilibration of small molecules and solute (*9*, *10*).

In this study, we characterized a novel semi-permeable capsule formulation composed of an acrylate-substituted polysaccharide polymer shell with a dextran core. Unlike conventional droplets, capsules are permeable, facilitating the selective exchange of biomolecules based on size. We demonstrate the utility of the capsule across diverse applications, including culturing clonally isolated cells, nucleic acid amplification, and conducting complex, multi-step molecular biology reactions such as single cell combinatorial indexing. Using these capsules we implemented a combinatorial indexing based single cell amplicon and single cell whole genome sequencing (sc-WGS) method capable of generating highly uniform single cell genomes at scale. Finally, to leverage the scale, coverage, and uniformity facilitated by capsule-based whole genome amplification and sequencing, we conducted a proof-of-concept lineage tracing experiment, using cells harboring a hypermutator polymerase ε allele that deposits mutations genome-wide with each cell division. These results, alongside two concurrent pre-prints (Baronas et. al. and Mazelis et. al.), highlight the array of single cell genomics applications that are uniquely enabled by capsules.

### Physical and chemical characterization of SPCs

Capsules are structures generated through the emulsification of two immiscible polymer solutions within droplets resulting in an aqueous two-phase system, where one polymer preferentially positions at the periphery of the droplet. The use of a shell monomer that is amenable to crosslinking, facilitates the formation of a hydrogel polymer shell. The shell polymer backbone can comprise synthetic polymers such as polyethylene glycol (PEG) (*9*) or natural polymers like gelatin (*10*). The choice of shell polymer fundamentally alters the properties of the capsules that are formed. For example, gelatin-based shells allow the controlled release of their content by mild enzymatic treatment, but this precludes the use of proteinases to process the content of the capsule, e.g. during cell lysis. Conversely, PEG-based SPCs are stable under a broader spectrum of treatment conditions but the release of their content requires mechanical or alkaline treatment destructive to cells (*9*). In this study we used a commercially available and novel Semi Permeable Capule (SPC) formulation consisting of an acrylate-substituted polysaccharide shell and a dextran core. As its predecessors (*9*, *10*), polysaccharide-based SPCs rely on droplet microfluidics for uniform high-throughput generation and sample encapsulation, followed by shell crosslinking and transfer into an aqueous suspension (**Figure 1A, Supplemental Video 1**). This creates compartments enabling the rapid exchange of buffers, and the diffusion of small reactants, while confining large reactants such as biological polymers, small solids and cells. The use of a polysaccharide-based shell also facilitates rapid release upon treatment with a glycosidase (**Supplemental Video 2**).

**Figure 1.**
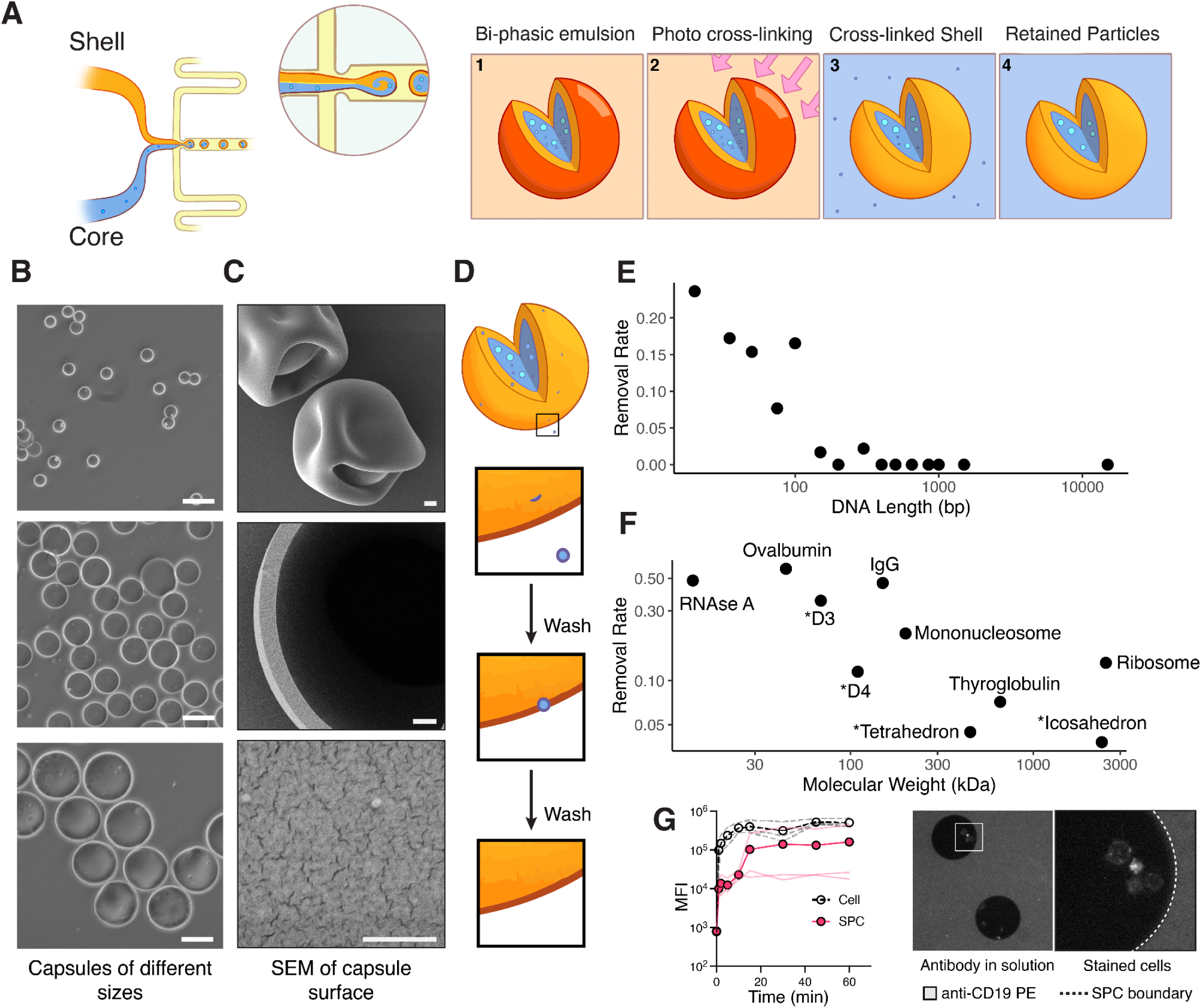
Semi-permeable capsules are tunable, porous containers. **(A)** Diagram of shell and core mixing upon encapsulation using a co-flow microfluidics device (left). Emulsions start in the oil phase, are cross-linked, and broken into the aqueous phase. Small molecules diffuse out of the porous shell (right). **(B)** Brightfield images of three capsule diameters (40.4µm (top), 82.9µm (middle), and 182.6µm (bottom). White scale bar represents 100µm. **(C)** SEM images of dehydrated capsules (scale bar 1µm), broken capsules (scale bar 1µm) and the surface of a capsule (scale bar 300nm). **(D-F)** Rate of removal was estimated by fitting an exponential loss function to retained biomolecules in SPCs after rounds of washing. (**E**) Removal rate for DNA species of different lengths, or (**F**) native and *de novo* designed (denoted by *) proteins by molecular weights. (**G**) (Left) Entry of antibodies into capsules over time as assessed by flow cytometry (red) versus staining of cells in solution (white). (Right) Representative image of encapsulated cells shown in a solution of fluorescently tagged antibodies (grayscale).

Monodisperse SPCs can be formed spanning a range of sizes from 40.4µm (SD = 1.97µm) in diameter to 182.6µm (SD = 16.3µm) in diameter by modifying the geometry of the microfluidics device used (**Figure 1B, Supplemental Table 1**). In this study, we primarily used SPCs formed with a mean diameter of 82.9µm (SD = 6.33µm). Remarkably, SPCs are stable, as evidenced by the compartmentalization of nucleic acids through overnight exposures to various organic solvents (e.g. hexane, methanol, ethyl acetate, and diethyl ether), strong acids and bases from 6M HCl to 6M NaOH, and resistant to dissolution from a battery of hydrolytic enzymes including nucleases and proteases (**Fig. S1**). The tolerance of the SPC shell to extreme physical and chemical conditions permits a wide array of manipulations on encapsulated material. In sum, SPCs are micron-sized containers that can be readily formed, highly stable, and can be broken on-demand.

**Supplementary Figure 1.**
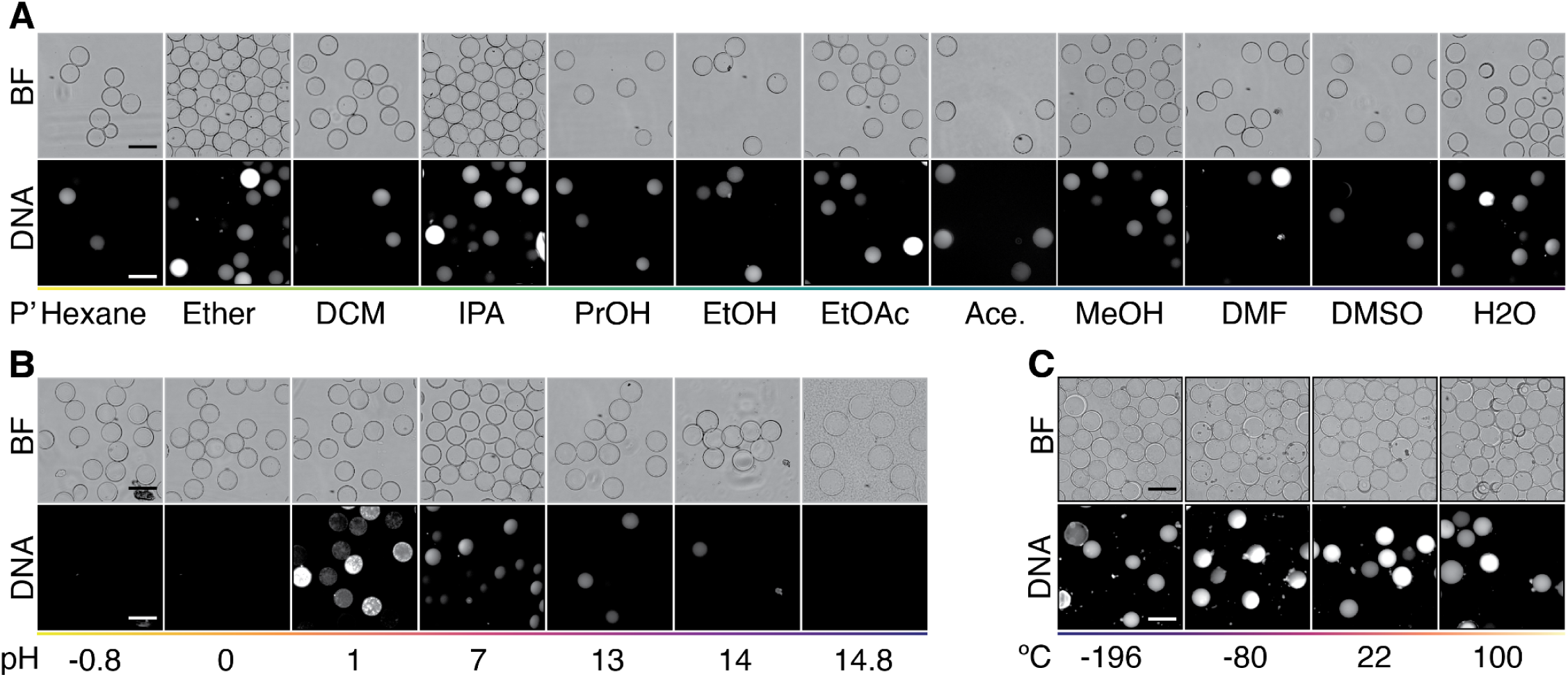
Capsule stability in extreme conditions. (**A-C**) Structural integrity testing of SPCs loaded with genomic DNA subject to overnight (>12 h) exposure to the indicated conditions. (**A**) Organic solvent exposures; Ether = diethyl ether, DCM = dichloromethane, IPA = isopropyl alcohol, PrOH = 1-propanol, EtOH = ethanol, Ace. = acetone, MeOH = methanol, DMF = N,N-dimethylformamide, DMSO = dimethyl sulfoxide. (**B**) Acid or base exposures; pH conditions represent 6M HCl (pH = -0.8), 1M HCl (pH = 0), 0.1M HCl (pH = 1), water (pH = 7), 0.1M NaOH (pH = 13), 1M NaOH (pH = 14), or 6M NaOH (pH = 14.8). (**C**) Temperature exposures; temperature conditions represent liquid N_2_ (-196°C), dry ice (-80°C), room temperature (22°C), or boiling water (100°C). BF = brightfield; DNA was stained with SYBR-Gold nucleic acid stain and representative images captured at 10× magnification; scale bar represents 50µM.

### Permeability of SPCs

To characterize the porosity of SPCs, we visualized SPCs by scanning electron microscopy (SEM). After dehydrating the sample, the exterior and interior surfaces of both intact and mechanically disrupted SPCs were visualized. At low magnification, the shell appeared to have a smooth interior and exterior surface with occasional irregularities. At higher magnifications (100,000x magnification), the shell polymer appeared to have a cracked structure (**Figure 1C**). To compare these results to hydrogels commonly used for electrophoresis, we prepared and imaged 6% polyacrylamide beads under identical conditions. Unlike the shell polymer, the surface of polyacrylamide was more mesh-like and dotted by irregularly spaced holes (**Fig. S2A)**.

**Supplementary Figure 2.**
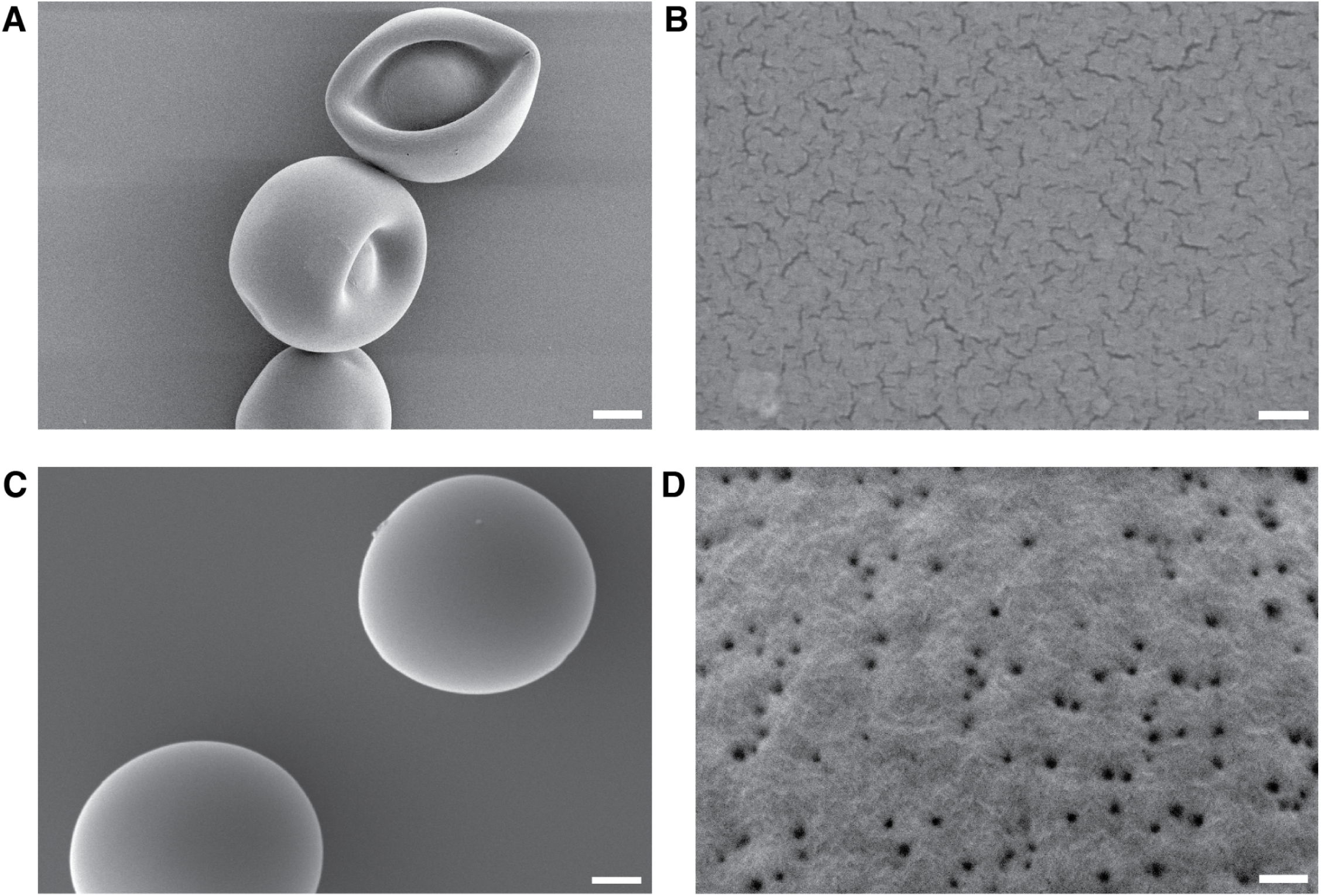
SEM comparison of semi-permeable capsules and polyacrylamide beads. (**A,B**) Micrographs of hexamethyldisilazane dehydrated (**A**) capsules at 10,000x magnification (scale bar 10µm) and (**B**) 100,000x magnification (scale bar 100nm). (**C,D**) Micrographs of HMDS dehydrated 6% polyacrylamide (PAA) beads at (**C**) 10,000x magnification (scale bar 1µm) and (**D**) 100,000x magnification, (scale bar 100nm).

Next we performed experiments to understand how biomolecules including DNA, protein, and nucleoprotein complexes were retained within SPCs. To this end, two DNA ladders spanning lengths over four orders of magnitude (10bp to 10,000bp), were loaded into SPCs. Fractions of SPCs were then collected in a serial manner, with washes performed between each collection over the course of 1 hour. Each SPC fraction was mechanically disrupted and the contents remaining within SPCs were quantified by gel electrophoresis. These experiments indicated that DNA fragments under 100bp diffused from the capsules and into solution over several washes. DNA with lengths ranging from 100 to 300bp, were lost at an average rate of 5.1% per wash, while larger fragments of DNA were retained completely (**Figure 1D, Fig. S3A**). We repeated this procedure using a native protein ladder, commonly used to calibrate size exclusion chromatography columns (**Fig. S3B**). This data revealed that the majority of encapsulated proteins below 300kD diffused out of the SPC over multiple washes (**Figure 1E**). To measure the retention of native cellular complexes, we purified and encapsulated mononucleosomes and ribosomes within SPCs, two complexes critical to the central dogma. Mononucleosomes and ribosomes were retained at 46.8% and 47.3% of the load after extensive washing, respectively (**Fig. S3C,D**). Unexpectedly, the ribosome was lost at rates exceeding the dimeric protein thyroglobulin despite having a greater molecular weight.

**Supplemental Figure 3.**
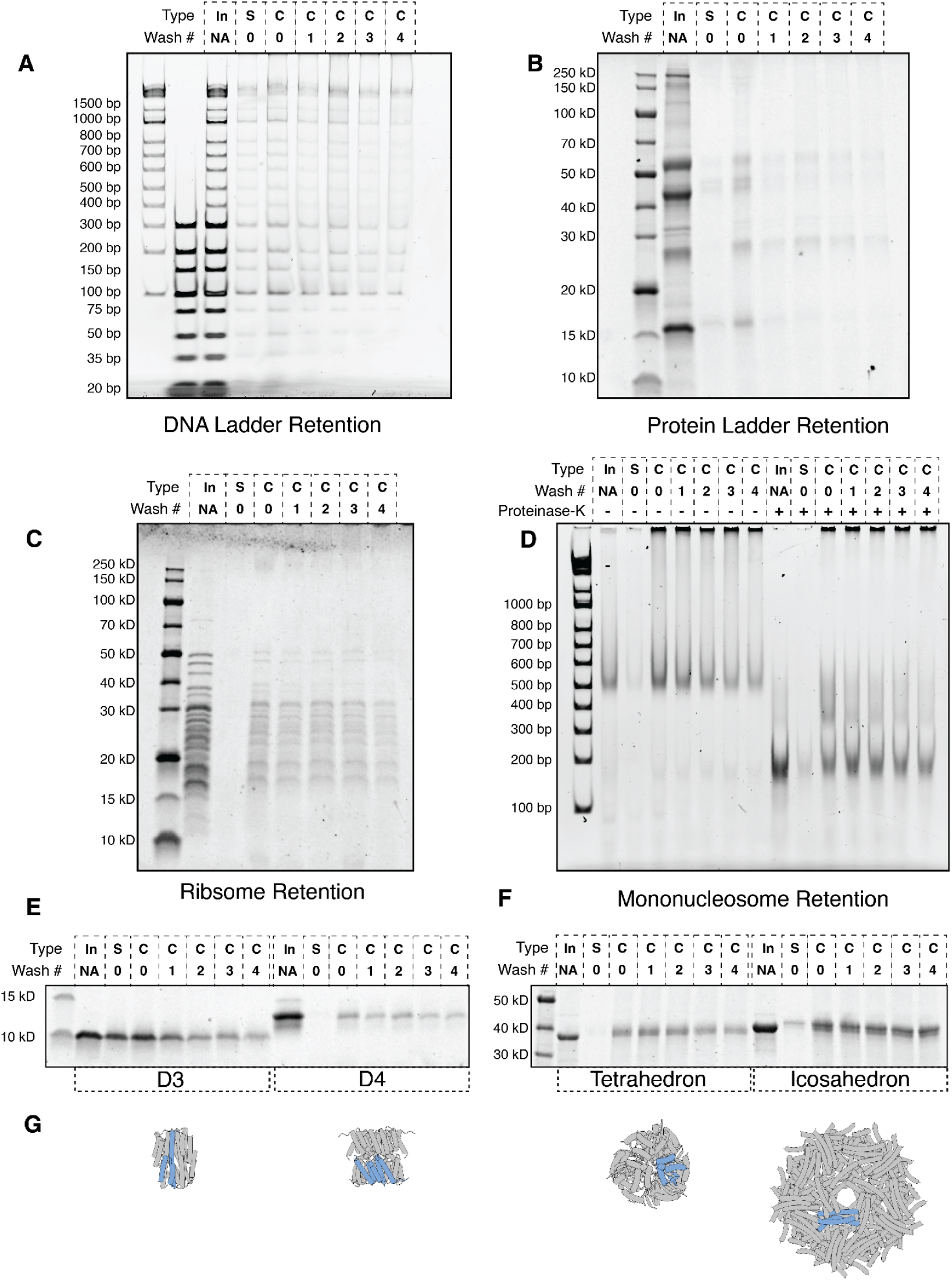
Retention of biomolecules within SPCs. SPCs loaded with **(A)** DNA ladder, **(B)** protein ladder, **(C)** ribosomes, **(D)** mononucleosomes, or **(E,F)** purified protein oligomers were broken after washing and the contents were run on gel electrophoresis and quantified. (**G**) Design models of the oligomers tested with a single monomer highlighted in blue. Legend for Type; In:Input. S:Supernatant, C:Capsule.

Many natural protein complexes have flexible three-dimensional structures and heterogeneous subunit interfaces spanning a range of affinities. Although these biomolecules are representative of cellular structures, these caveats preclude their use as molecular calipers for the determination of the shell’s porosity. To overcome this limitation, we used *de novo* designed proteins to generate a set of rigid and atomically-precise oligomers spanning various symmetry point groups. Three published *de novo* designed oligomers (*11*) were chosen in addition to a newly designed oligomer for retention testing. These proteins included a D3 (6-mer), a D4 (8-mer), a tetrahedron (12-mer) and an icosahedron (60-mer), spanning a range of molecular weights (**Fig. S3E-G**). When encapsulated in SPCs, purified *de novo* designed oligomers exhibited retention within SPCs in a size-dependent manner (**Figure 1E**), and were stably retained within SPCs over the course of 10 days (**Fig. S4**).

**Supplemental Figure 4.**
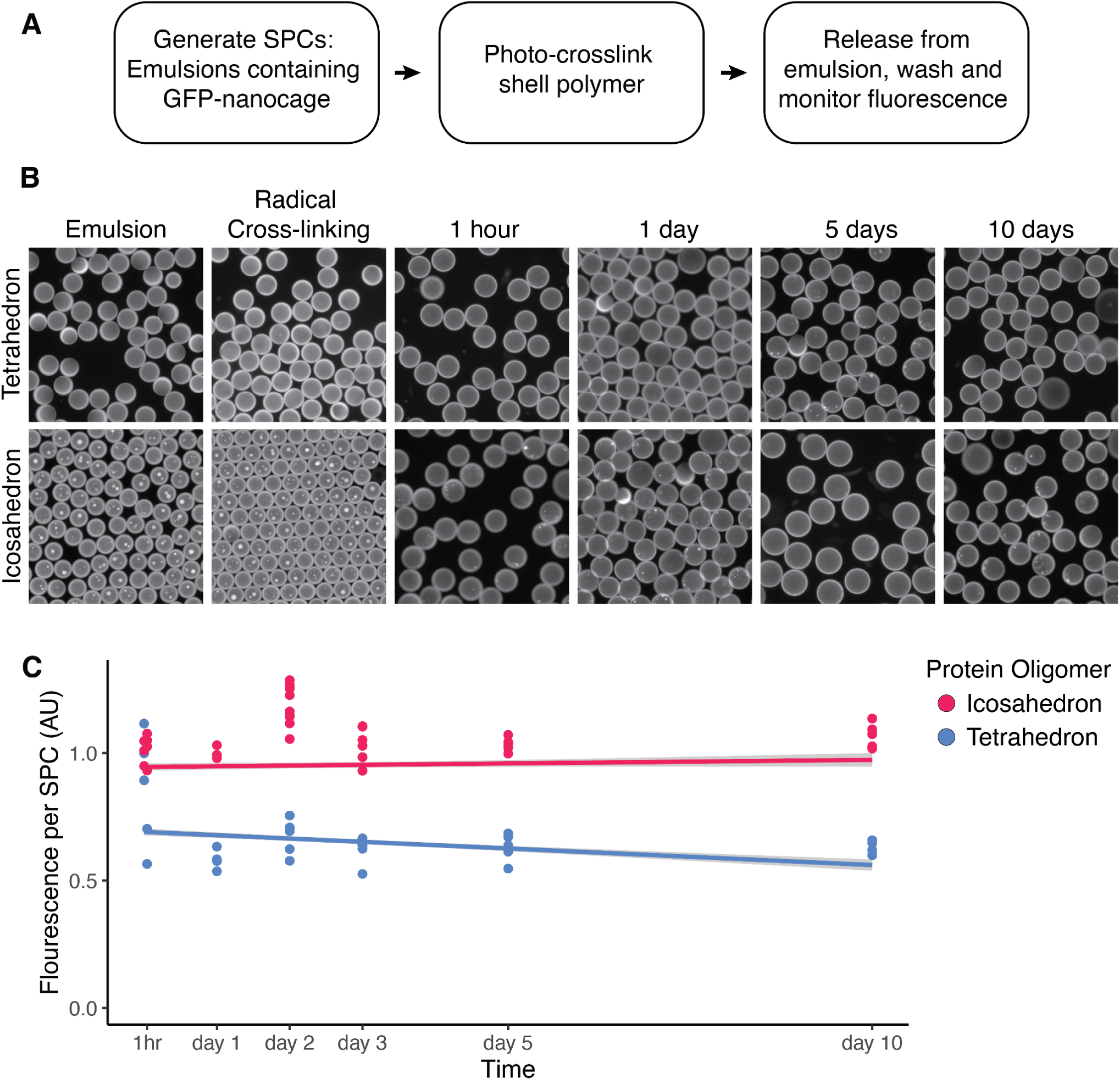
Protein oligomers are stably retained for many days. (**A**) Tetrahedron and icosahedron protein oligomers fused to GFP were expressed and encapsulated within SPCs. (**B**) Emulsions were imaged before and after cross-linking of the shell. Capsules were imaged at time intervals spanning 10 days using the same imaging settings. (**C**) Quantification of the fluorescence within each SPC was quantified from multiple images. Points represent the mean cumulative fluorescence of volumes within each image and the line depicts the linear best fit with standard error shown in gray.

The diffusion of biomolecules across a concentration gradient can be used to introduce molecules into SPCs as well. To demonstrate this, we attempted to fluorescently stain cells after they had been encapsulated within SPCs. We incubated SPCs containing Nalm6 cells, a leukemic B cell line that expresses high levels of the surface marker CD19, in a solution containing a phycoerythrin-conjugated anti-CD19 antibody (anti-CD19 PE). Using flow cytometry on released cells, our experiments demonstrated that IgG diffused across the SPC membrane and bound to the surface of encapsulated cells mirroring the diffusion of IgG out of SPCs (**Figure 1G**, **Fig. S5**). In sum, these data indicate that SPCs are permeable to short oligonucleotides (e.g. primers and small double stranded DNA) and proteins (e.g. enzymes and IgGs), and selectively retain larger nucleoprotein complexes including chromatin and ribosomes. These permeability characteristics of SPCs make them an ideal container for many cellular and molecular assays.

**Supplementary Figure 5.**
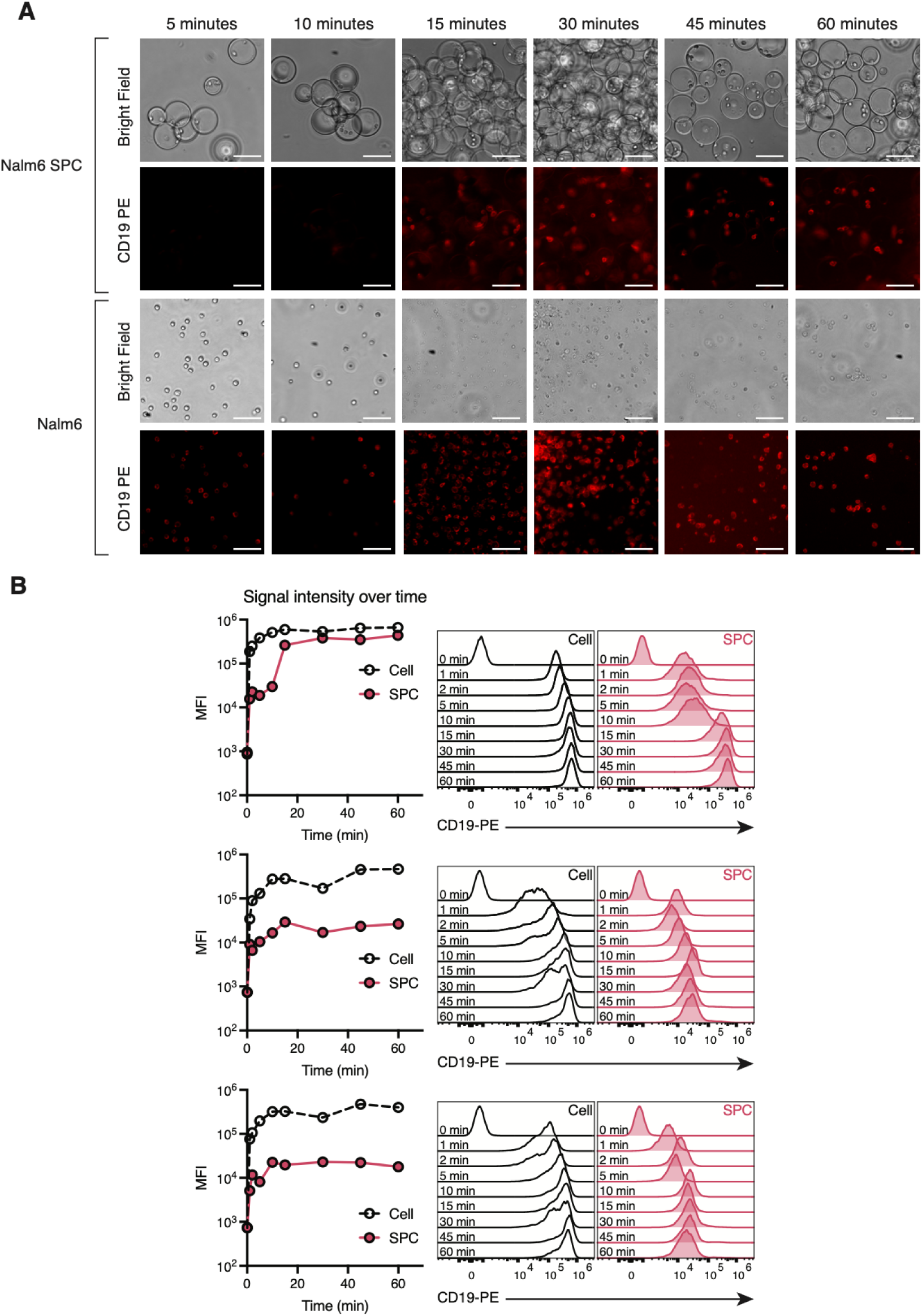
Fluorescent staining in semi-permeable capsules. (**A**) Nalm6 were encapsulated in SPCs and then stained for different incubation times with a fluorescent antibody targeting CD19 (CD19-PE) displayed on the surface of the cells and compared to free Nalm6. Brightfield (top) and fluorescent (bottom) images are shown for incubation times of 5, 10, 15, 30, 45, and 60 minutes for encapsulated Nalm6 (top set) and free Nalm6 (bottom set) (scale bar 75µm). (**B**). Flow cytometry quantification of fluorescence signal for each experimental replicate. Median fluorescence intensity over time (left) and corresponding flow plots (right).

### High throughput cell culture within SPCs

Emulsified cells within microfluidic droplets are viable and can be grown for days. However, due to compartmentalization of emulsions, cells exhaust nutrients needed for anabolism, while accumulating the toxic byproducts of cellular metabolism (*7*, *8*). Moreover, upon the dissolution of an emulsion, the physical link between clonally expanded cells is lost. We hypothesized that the permeability of SPCs to small molecules and small proteins would facilitate the exchange of nutrients to support the long term culture of bacterial and mammalian cells within SPCs at scale. To test this hypothesis, *E. Coli* were encapsulated within SPCs at a poisson loading rate (21% occupancy rate – 525,000 loaded SPCs; 2.37% multiplet rate) and grown in a shaking culture. After measurement, *E. Coli* grown in SPCs had a doubling time of approximately 20 minutes, comparable to the doubling time of 29 minutes for bacteria cultured in suspension (**Figure 2B**). To test if clonal bacterial isolates could grow within SPCs, a 50:50 mix of bacteria expressing one of two fluorescent markers (mCherry or eGFP) were mixed prior to encapsulation, imaged, and analysed after bacterial growth. If cells grow in a clonal fashion, we expect to predominantly observe capsules marked by a single fluorescent bacterial species and a small population of capsules containing both species. Indeed, in accordance with the poisson loading rate, we observed SPCs that were either empty (71.6%), mScarlet positive (11.3%) or eGFP positive (13.8%), with a small portion of SPCs (3.2%) exhibiting a mixed population of cells indicating that we could establish clonal bacterial cultures within SPCs (**Figure 2C,D**).

**Figure 2.**
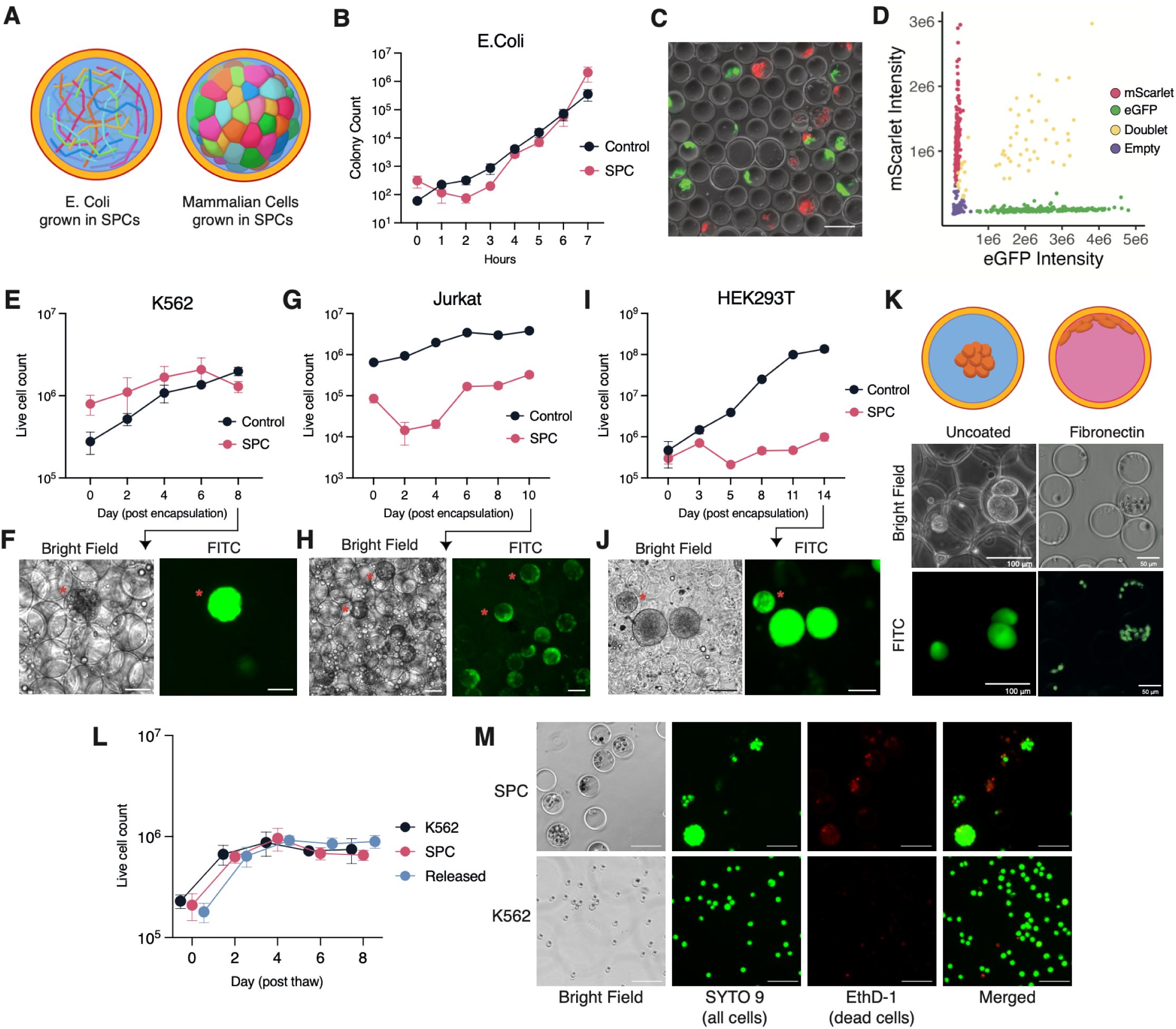
Semi-permeable capsules facilitate scaled and compartmentalized cell culture. (**A**) Schematic depicting E.Coli or mammalian cells encapsulated and grown within SPCs. (**B**) Growth curves for *E. Coli* in suspension (black) or grown within SPCs (red). (**C**) Micrograph of SPCs poisson loaded with a 50:50 mix of eGFP or mScarlet expressing bacteria and (**D**) quantification of total fluorescence from each segmented SPC. (**E**-**J**) Growth curves for fluorescent mammalian cell lines and images for (**E,F**) K562, (**G,H**) Jurkat cells and (**I,J**) HEK293T cells. Red asterisks denote select SPCs positive for mammalian cells (**K**) Image of HEK293T cells grown in untreated (left) or fibronectin treated (right) SPCs. (**L**) Growth curves for K562 cells frozen in SPCs and thawed SPCs (red) or thawed and released (blue) versus cells frozen in suspension (black). (**M**) Representative images of cell viability two days after freezing and thawing in SPCs. Scale bars represent 50µm (F,H) and 100µm (J, C, M)

Next we tested the viability and growth rates of common suspension and adherent mammalian cell lines within SPCs. The growth of a cell suspended within the SPC’s core solution most closely resembles the growth of cells suspended in liquid media. Accordingly, we used an eGFP-expressing suspension cell line (K562 cells) to assess the viability and growth rates of mammalian cells within SPCs. After cell encapsulation (44.5% occupancy rate – 1,112,500 loaded SPCs; 11.8% multiplet rate) and subsequent culture, we found that K562 cells were not viable within SPCs. By day 4 post encapsulation, cell viability had decreased by approximately 72%, and by day 8 nearly all the cells were dead (**Fig. S6A**). Given our previous observation that diffusion rates across the SPC shell are diminished, we hypothesized that the limited diffusion of nutrients and signaling factors created an environment unsuitable for single-cell growth. To test this hypothesis, we repeated this assay targeting a higher cell encapsulation density within SPCs (69.16% occupancy rate – 1,729,000 loaded SPCs; 32.9% multiplet rate) and substituted fresh media with conditioned media. Under these new culture conditions, encapsulated K562s exhibited sustained growth (**Figure 2E,F**). Based on imaging and segmentation, we estimate that up to approximately 60 K562 cells are able to grow within the capsule (**Fig. S6H,I**) until physical crowding prevents further cell divisions (**Fig S7A**).

**Supplementary Figure 6.**
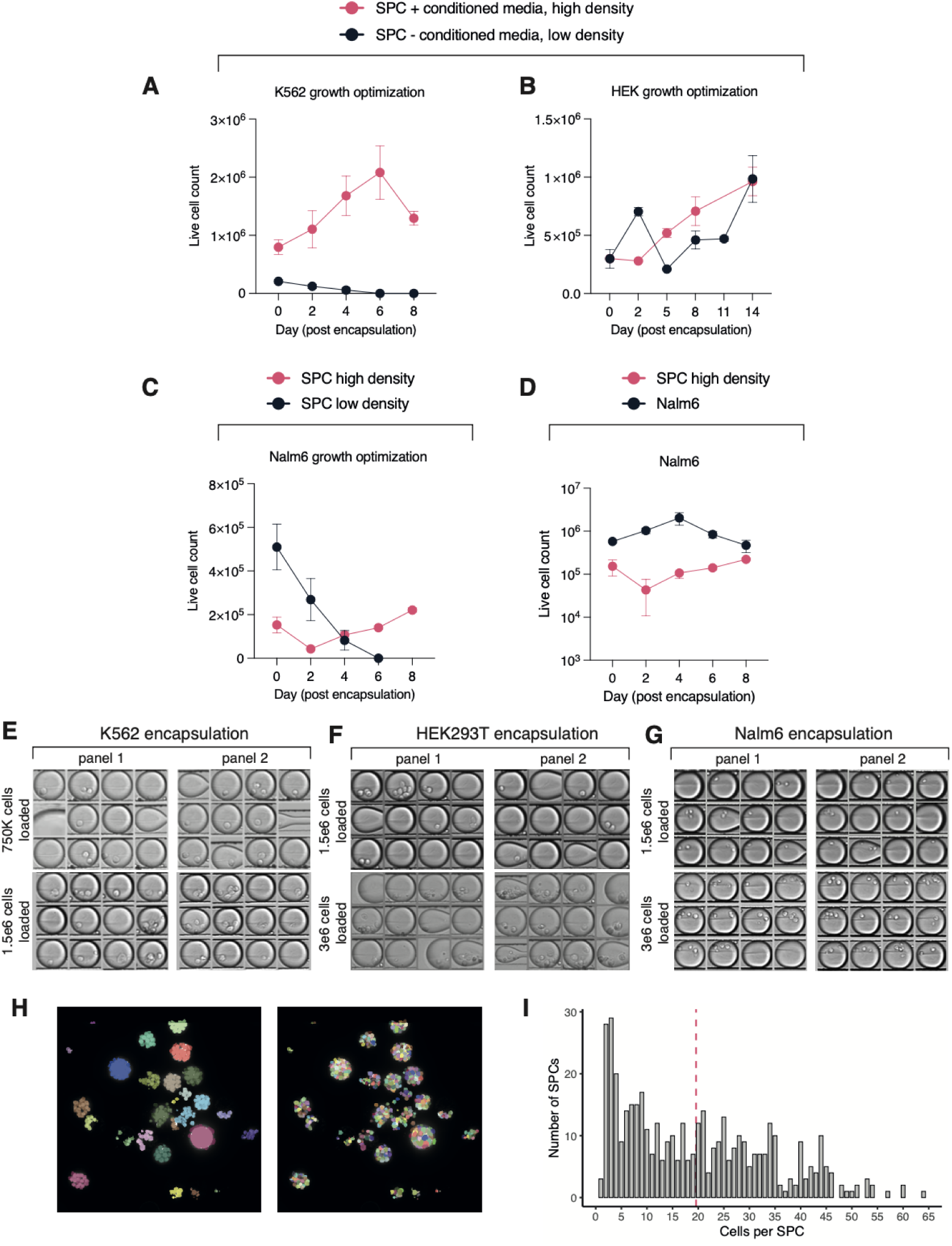
Mammalian growth optimization in SPCs. (**A**) K562 were encapsulated in fresh media at low density (7.5× 10^5^ cells) or in conditioned media at high density (1.5 × 10⁶ cells). (**B**) HEK293T were encapsulated in conditioned media at low density (1.5 × 10⁶ cells) or in conditioned media at high density (3× 10⁶ cells). (**C**) Nalm6 cells were encapsulated in conditioned media at either low density (1.5 × 10⁶ cells) or high density (3 × 10⁶ cells). (**D**) Comparisons of encapsulated Nalm6 cell growth rate with non-encapsulated control cells. (**E–F**) Representative crops of encapsulated cells at high and low densities, corresponding to panels A–C. (**H**) (Left): Individual K562 SPCs were stained with Yo-Pro-1 DNA dye and imaged with a spinning-disk confocal microscope, 60x magnification. Colors represent individual SPCs, assigned by identifying clusters of nuclei segmentation masks with DBSCAN. (Right): Single-nucleus segmentation with CellPose 3.0 ’cyto3’ base model. **(I)** Histogram of K562 cells per SPC, computed by counting nuclear segmentation masks assigned to each SPC by DBSCAN. Dashed red line marks the median of 20 cells per SPC.

**Supplementary Figure 7.**
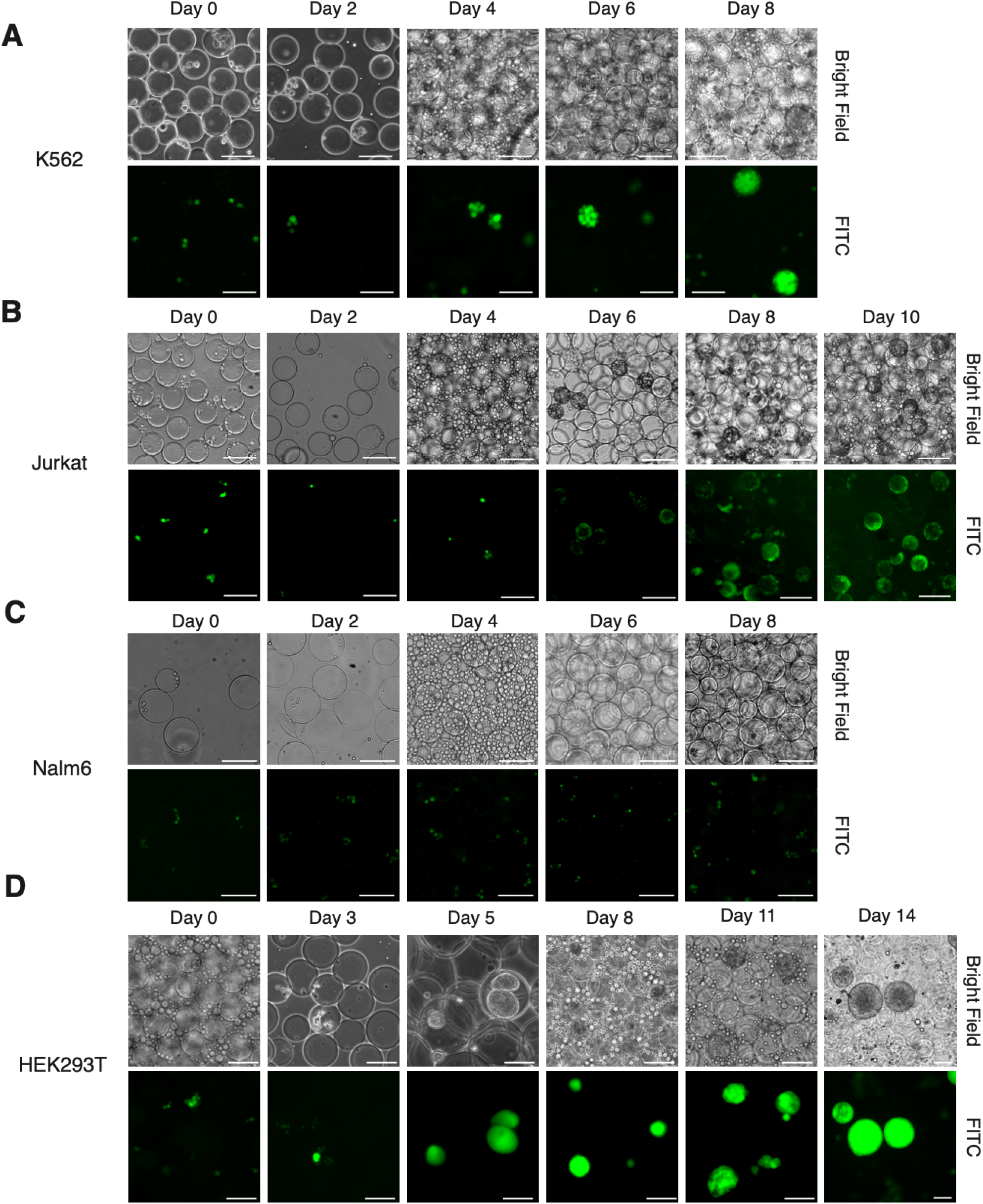
Cell growth time course in SPCs. Brightfield (top row) and fluorescent (bottom row) images representative of mammalian cell growth within SPCs for (**A**) K562, (**B**) Jurkat, (**C**) Nalm6 and (**D**) HEK293T cells taken regularly between an eight and fourteen day interval. Scale bars represent 100µm.

To assess the generalizability of these findings to other cell lines, we prepared capsules containing additional suspension cell lines (Nalm6, Jurkat) or an adherent cell line (HEK293T) (**Fig. S7B-D**). For all three cell lines we observed an initial decrease in cell count, followed by sustained cell growth (**Figure 2G-J, Fig. S7B-D**). We suspect that this decline reflects the death of single cells isolated within SPCs and the selection for cells within SPCs containing multiple cells. Notably, adherent HEK293T cells adopted a distinct spheroid morphology as early as day 5 (**Figure 2K**), forming a clear separation between the cell mass and the polymer shell wall. Despite a lack of adhesion to the polymer wall, HEK293T cells continued to divide, forming spheroids that expanded to fill the SPCs and stretching the capsule beyond its typical size (**Fig. S7D**). Notably, this behavior could be modified upon co-encapsulation with fibronectin, effectively tissue culture treating the SPC shell polymer and causing HEK293T cells to grow along the boundary of the capsule (**Figure 2K**). Finally, we assessed whether cellular colonies grown in capsules could be frozen and cryo-recovered within capsules. To this end, we cryopreserved K562 cells in SPCs, and subsequently thawed these cells either as capsules or in the presence of a release agent. These results showed no significant difference in viability or growth rates between cells frozen in suspension or cells thawed in SPCs. (**Figure 2L,M**, **Fig. S8,9**). Together, these experiments demonstrate the ability of SPCs to support clonally isolated cellular cultures, and highlight their potential as a novel platform for the generation and long-term storage of 3D cell cultures.

**Supplementary Figure 8.**
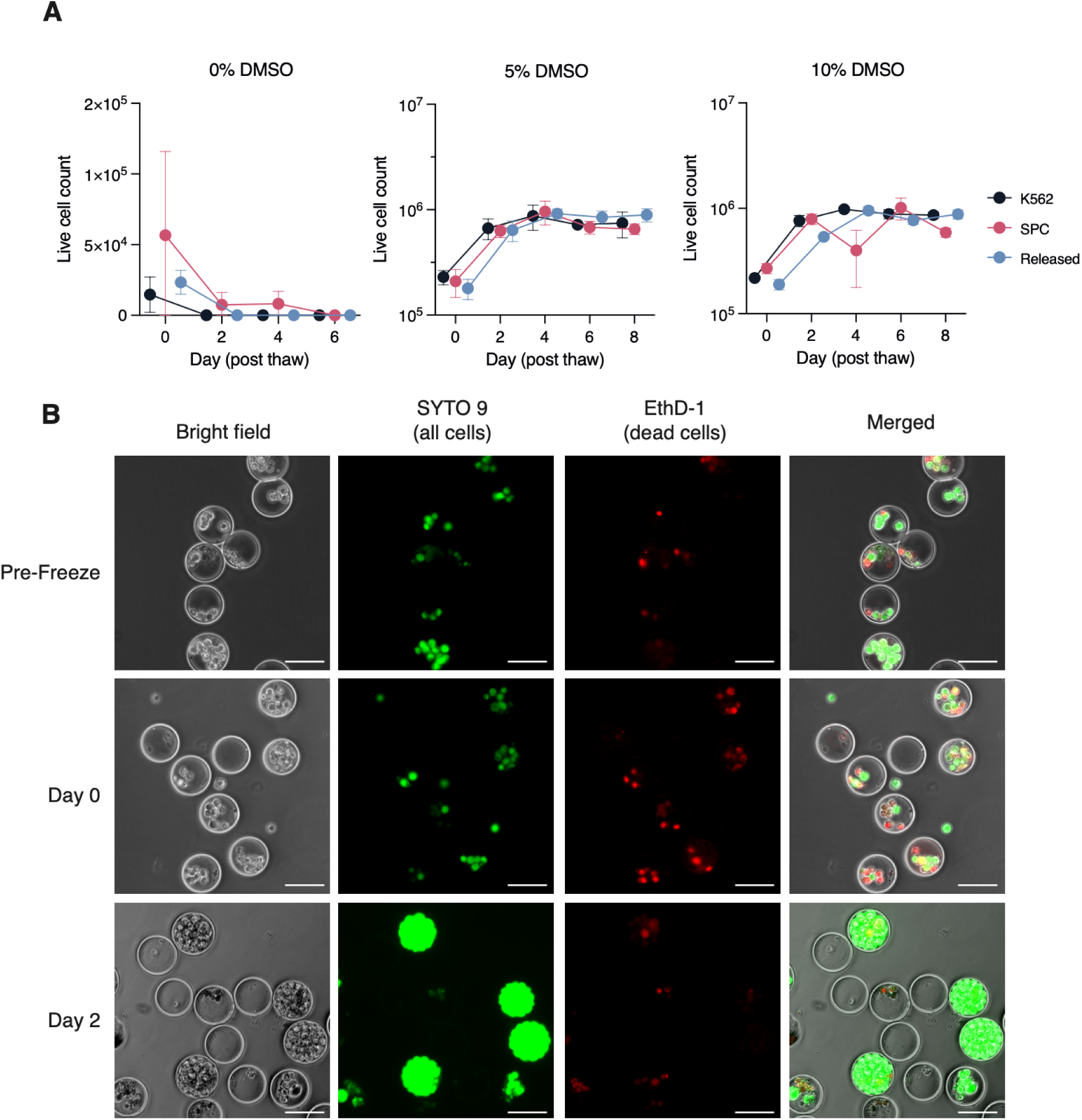
K562s can be frozen and thawed in SPCs. (**A**) Live cell counts over time of thawed cells frozen under DMSO concentrations of 0% (left), 5% (middle), and 10% (right). (**B**) Live/dead imaging of cells encapsulated in SPCs before freezing in 10% DMSO media (top), immediately after thawing (middle), and 2 days after thawing (bottom). Scale bars represent 75µm.

**Supplementary Figure 9.**
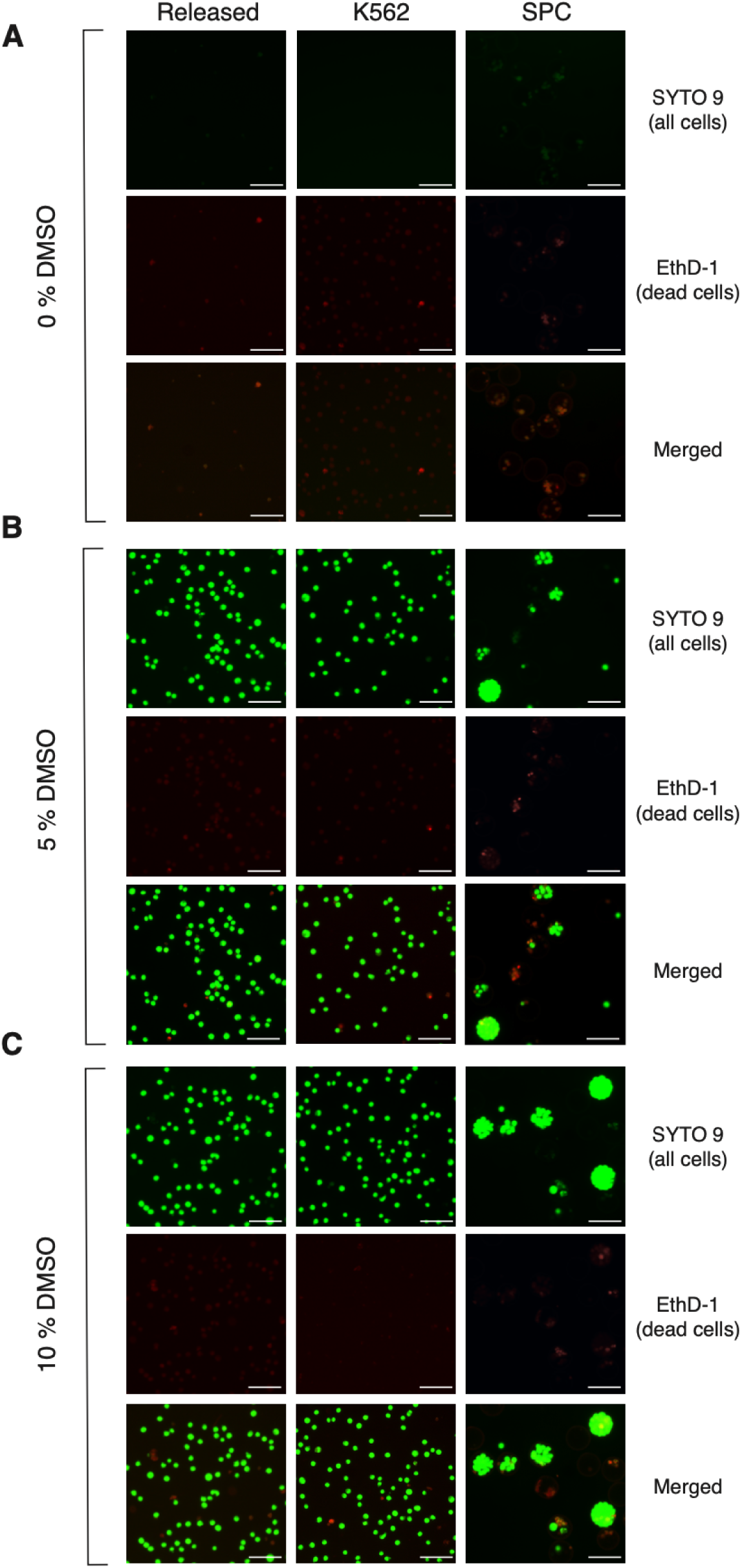
K562 freeze-thaw comparison Day 2. (A-C) K562 were encapsulated at a initial cell density 3 × 10⁶ cells and were frozen in one of three media, all consisting of DMEM, 10% FBS, and a varying percent of DMSO: 0%, 5%, 10%. Images were taken two days post thawing, after staining cells with SYTO 9 and EthD-1 to assess viability. Scale bars represent 100µm.

### SPC Polymerase Chain Reaction

The retention of nucleic acids within capsules has been used previously for applications relating to the amplification and detection of DNA sequences from single cells (*10*). To further characterize SPCs comprising a polysaccharide shell, we tested the performance of nucleic acid amplification within SPCs (**Figure 3A**). First cells were encapsulated, permeabilized, and digested to remove lipids and proteins. Capsules, containing clarified genomic DNA (gDNA), were then used as the template for polymerase chain reaction (PCR), where enzymes, primers, and cofactors were introduced outside of the capsule. We anticipated that smaller DNA amplicons would diffuse out of the capsule based on the retention tests performed for double stranded DNA. To test the relationship between amplicon length and retention within capsules during PCR, we generated primers spanning a range of amplicon sizes from 100bp to 1,500bp in length. After PCR, the molecules retained within SPCs and the molecules in solution were analyzed by gel electrophoresis. Strikingly, we observed amplicons within SPCs and in solution for all the amplicon sizes (**Fig. S10**). To test whether this was a function of thermocycling during PCR, we also tested the retention of DNA during isothermal whole genome amplification using primary template amplification (PTA), a reaction performed at 30°C. Analysis of the supernatant and capsule fractions after amplification showed the presence of DNA ranging from 150bp to 1,500bp in the supernatant indicating the leakage of amplified template even in the absence of thermocycling and elevated temperatures (**Fig. S11**). Although nucleic acid amplified outside of capsules can be removed by washing, the entry of amplicons from the supernatant and into another SPC would be problematic for single cell assays. To test the prevalence of amplicon entry during PCR, we monitored the entry of amplicons into empty capsules during a PCR performed in the supernatant. Amplicons of different lengths were subject to 30 rounds of PCR in the supernatant in the presence of empty capsules. Capsules from each PCR reaction were then washed extensively and subjected to a second round of PCR to amplify material that may have entered from the supernatant. For amplicons ranging in size from 100bp to 500bp we observed a faint band at the expected size indicating that some amplicons had indeed entered the empty capsule during the first round of PCR (**Fig. S12**). Although these bands were present after two rounds of PCR, visual examination of individual SPCs upon DNA staining indicated that only a subset of the SPCs contained DNA.

**Figure 3.**
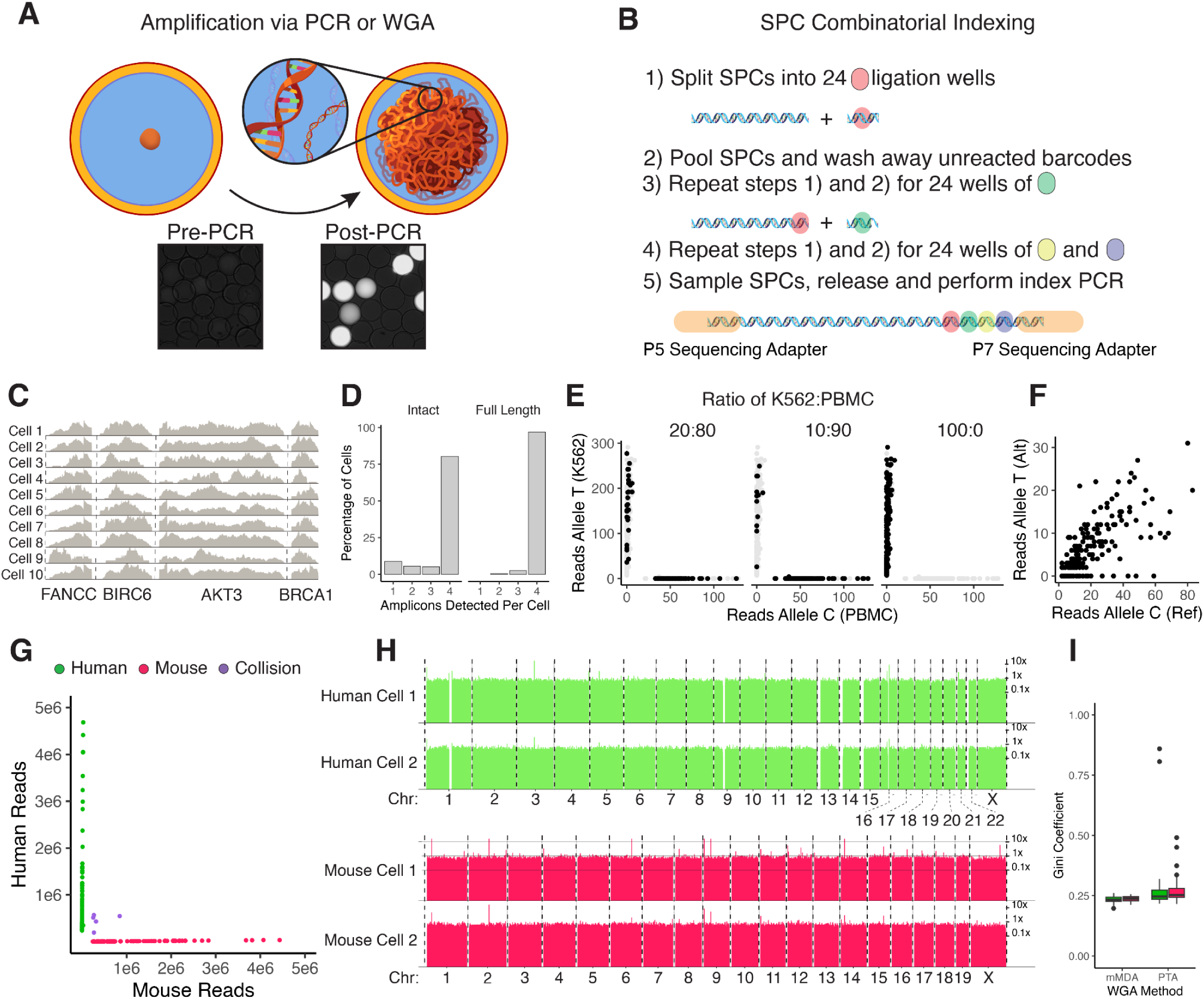
Single cell DNA sequencing via combinatorial indexing in SPCs. (**A**) Nucleic acids within SPCs are readily amplified via PCR or whole genome amplification (WGA). Fluorescent microscopy image of DNA stained SPCs is shown before amplification (bottom left) and after amplification (bottom right). (**B**) Diagram describing combinatorial indexing procedure. (**C**) Coverage of 10 randomly selected single cell libraries mapping to the 4 amplified gene fragments. (**D**) Barplot showing the percentage of cells with the detection of the different amplicons in fragmented or intact amplicon libraries. (**E**) Faceted scatterplot showing the reads mapping to the *FANCC* gene position chr9:95,247,502 with either the reference allele (X–axis) versus the alternate allele (Y-axis). (**F**) Scatterplot for reads mapping to a heterozygous position chr1:243,695,654 in the *AKT3* gene for each cell. (**G**) Scatterplot of the number of reads uniquely mapping to the human (y-axis) or mouse (x-axis) genomes for each individual cell. (**H**) Coverage of cells amplified via PTA: Human genome (green, top) or Mouse genome (red, bottom). Genome binned at 1kB. (**I**) Boxplot showing Gini coefficient for cells amplified with either a modified MDA protocol (mMDA) or PTA.

**Supplemental Figure 10.**
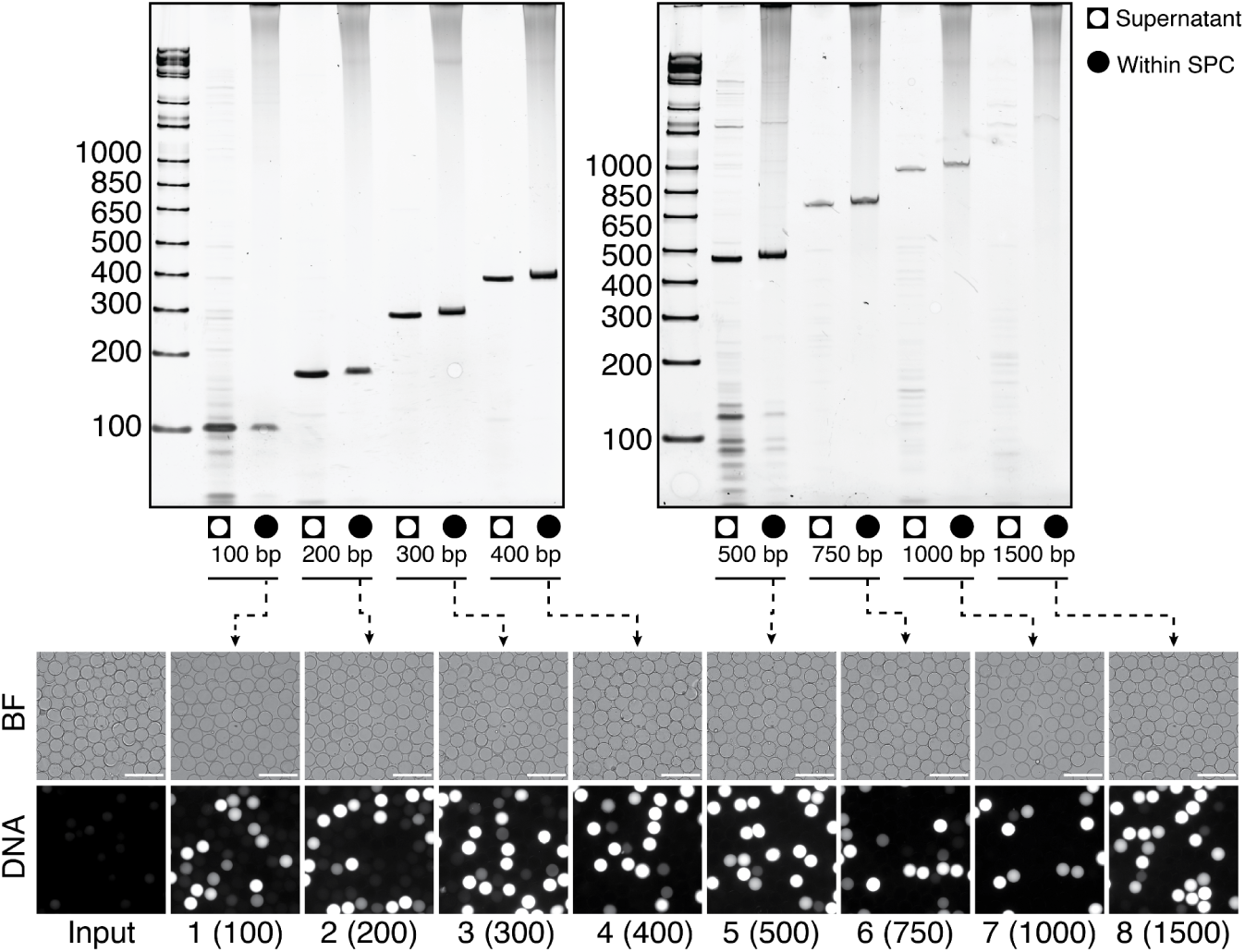
Amplicons leak out of capsules during PCR. Genomic DNA derived from cells within capsules was used as the template for PCR. Primers spanning a single genomic DNA locus were designed to capture amplicons of approximately 100, 200, 300, 400, 500, 750, 1000 and 1500 base pairs in length. Gel electrophoresis was performed on supernatant or washed and released SPCs (within SPC). Representative images of SPCs in bright field (BF) or stained with the DNA dye SYBR green (DNA) are shown below. Scale bars represent 500µm.

**Supplemental Figure 11.**
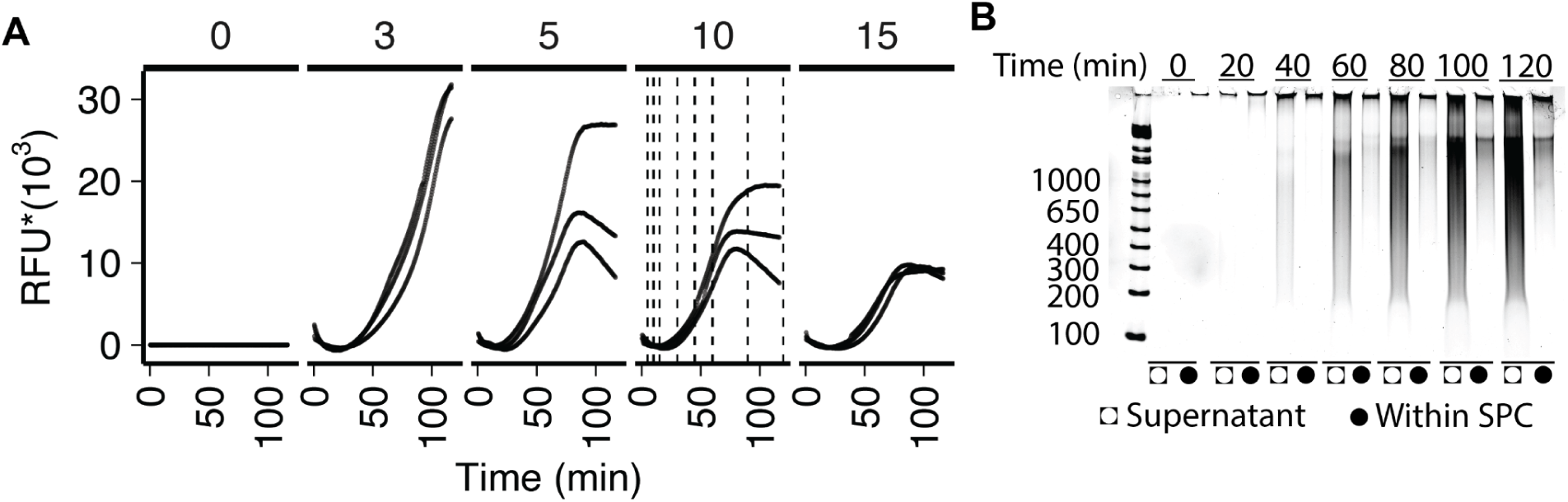
Genomic DNA originating in capsules is amplified in SPCs and the supernatant. (**A**) Titered volumes of SPCs containing genomic DNA were provided as input (0, 3, 5, 10 or 15μL) with fixed amounts of primary template amplification primers (thiophosphate and phosphodiester hexamer), dNTPs, and reaction cofactors. Relative fluorescence units (RFU) of SYBR-Green quantify total yield of DNA over real time (N=3 technical replicates). (**B**) From the 10μL condition in (A), supernatant and DNA retained within SPCs were collected at the indicated time points and run on a 1% TBE-Agarose gel and stained with SYBR-Gold.

**Supplemental Figure 12.**
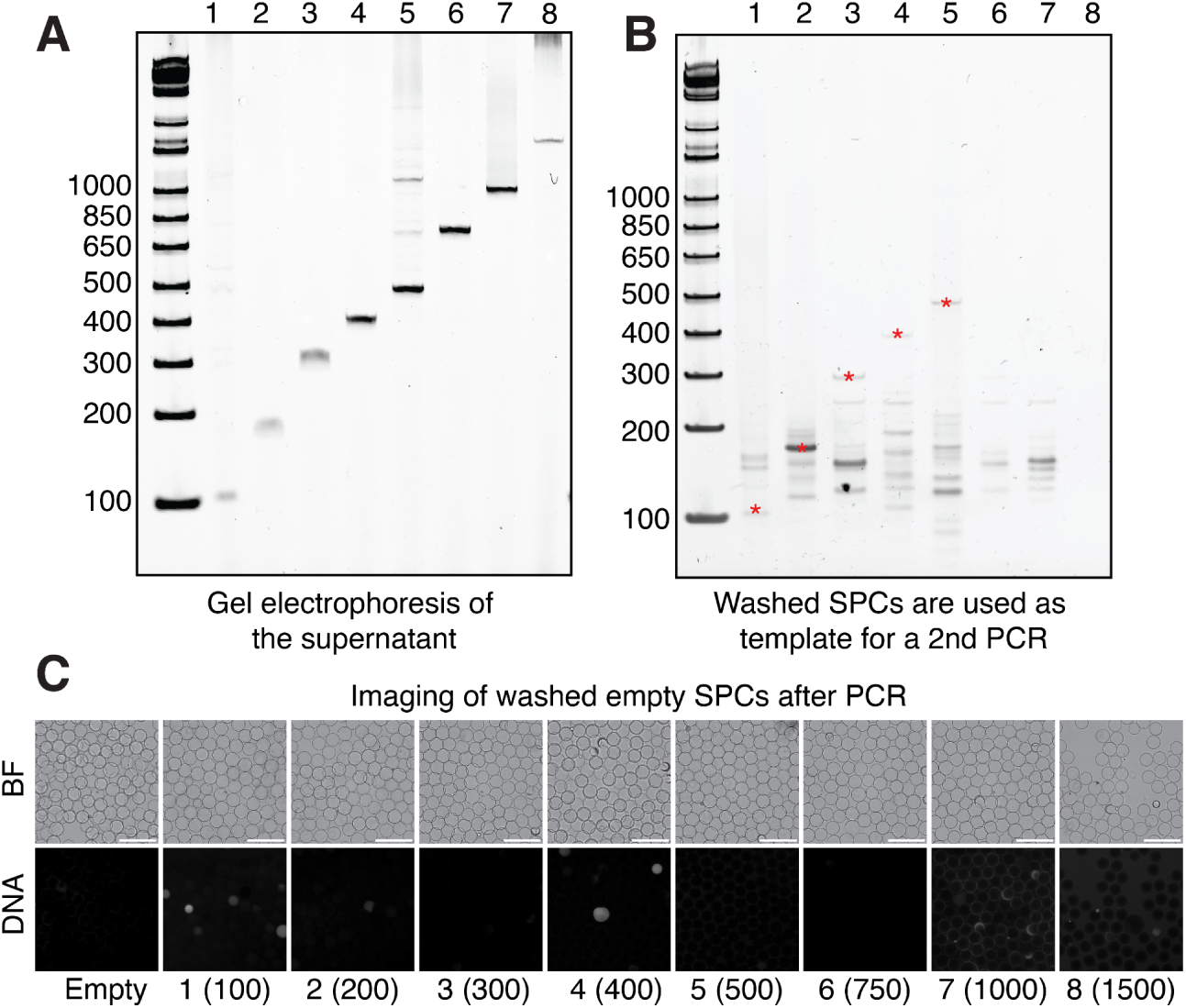
Amplicons leak into capsules during PCR. (**A**) PCR was performed with the template, primers and mastermix in the bulk in the presence of empty SPCs. (**B**) Nested PCR was performed after washing capsules using primers using the primers used during the first PCR. Gel denotes the products retained within capsules after nested PCR. Red stars denote the detection of a band of the expected size. (**C**) Representative brightfield and SYBR green stained images of capsules for each amplicon. Scale bars represent 500µm.

### Sensitive genome sequencing with SPC combinatorial indexing

The properties explored thus far make SPCs an ideal container for combinatorial indexing based single cell sequencing (*12–14*). Current combinatorial indexing methods are limited primarily by the stability of chemically fixed cells or nuclei to harsh detergent, elevated temperature, or enzymatic treatment (e.g. Proteinase K digestion) (*15*). Paradoxically, the perfect container for combinatorial indexing must simultaneously allow for the free diffusion of small biomolecules needed for barcoding (e.g. primers, and enzymes), while preventing the escape of nucleic acids. Guided by the principles gleaned from biomolecule retention and DNA amplification experiments, we devised a ligation-based combinatorial indexing protocol (*16*), where genomic DNA is pre-amplified within SPCs and then split-pooled to combinatorially synthesize unique cellular barcodes on all DNA species within SPCs (**Figure 3B**). To test whether amplicons originating from one SPC contaminate other SPCs, we encapsulated K562s or peripheral blood mononuclear cells (PBMCs) separately, and then mixed capsules at defined ratios (100% K562, 20:80 K562:PBMC, or 10:90 K562:PBMC) prior to multiplex PCR for 4 genes (*AKT3*, *BIRC6, FANCC and BRCA1*). Combinatorial indexing was then performed both on intact amplicons, as well as fragmented amplicons. All amplicons tested exceeded 400bp in length ranging from 460bp to 2,155bp in length (**Figure 3C**, **Supplemental Table 2**). Upon sequencing the full length amplicon library, we recovered 387 cells sequenced to a depth of 1,008 reads/cell and detected amplicons mapping to all 4 sites in 96.9% of the cells (**Figure 3D**). To assess whether mixing was occurring between cells, we investigated a locus in the FANCC gene (chr9:95,247,502) which is homozygous for an alternate allele (T) in K562s versus homozygous for the reference allele (C) in PBMCs. Cells with reads mapping to this allele segregated to the axes, indicating a minimal mixture of amplicons originating from this gene (**Figure 3E,F**). Closer investigation of cells prepared from the 10:90 and 20:80 mixtures of K562s:PBMCs – where an increased proportion of swapping would be expected – demonstrated that an average of 99.2% of reads (in 39 cells) mapped to the alternate allele in the intact amplicon libraries, while 100% of reads (in 20 cells) mapped to the correct genotype in fragmented amplicons. Finally, to determine the sensitivity to retrieve alleles within a single cell, we examined a locus in *AKT3* (chr1:243,695,654) in K562 cells that contains a heterozygous premature termination allele (p.G37Ter) (*17*). In the intact amplicon library we observed both alleles at this site in 84.91% of cells (mean of 28.34 reads/cell) (**Figure 3G**).

Encouraged by these results, we turned to the generation of sc-WGS libraries within SPCs. Whole genome amplification (WGA) methods often suffer from uneven or biased amplification (*18*, *19*). This bias is pronounced in sc-WGS, where comparisons between cells require observing the same genomic position in multiple cells. New molecular biology methods, which operate chiefly through the addition of chain terminating dNTPs, have largely mitigated this problem (*20*); however, these methods have limited throughput due to their reliance on well-based amplification (*21*). To perform scaled WGA and sequencing we used a commercial WGA reaction mix (Bioskryb PTA) or a modified multiple displacement amplification (mMDA) reaction to prepare sc-WGS libraries from a 50:50 mixture of encapsulated human and mouse cells. Following amplification, SPCs containing amplified genomes were combinatorially indexed, fragmented, and sequenced. After shallow sequencing (mean 1,433,349 reads/cell) and mapping to a jointly indexed genome, we observed that the reads from cells mapped predominantly to either the human or mouse genome, indicating minimal cross contamination during amplification and library preparation (**Figure 3G)**. Visual Inspection of genome coverage in individual cells indicated uniform coverage of the genome by both amplification methods (**Figure 3H**, **Fig. S13**). To quantify coverage uniformity, we generated Lorenz curves and computed Gini coefficients for both methods by binning the genome into 1kB bins and counting the number of reads within each bin. Both WGA procedures produced genomes with comparable and even coverage with average Gini coefficients of 0.236 for mMDA and 0.277 for PTA (**Figure 3I**). Together, these results demonstrate that both WGA approaches (mMDA and PTA) enable scalable and uniform amplification of single-cell genomes within SPCs with minimal cross-contamination during amplification or combinatorial indexing.

**Supplemental Figure 13.**
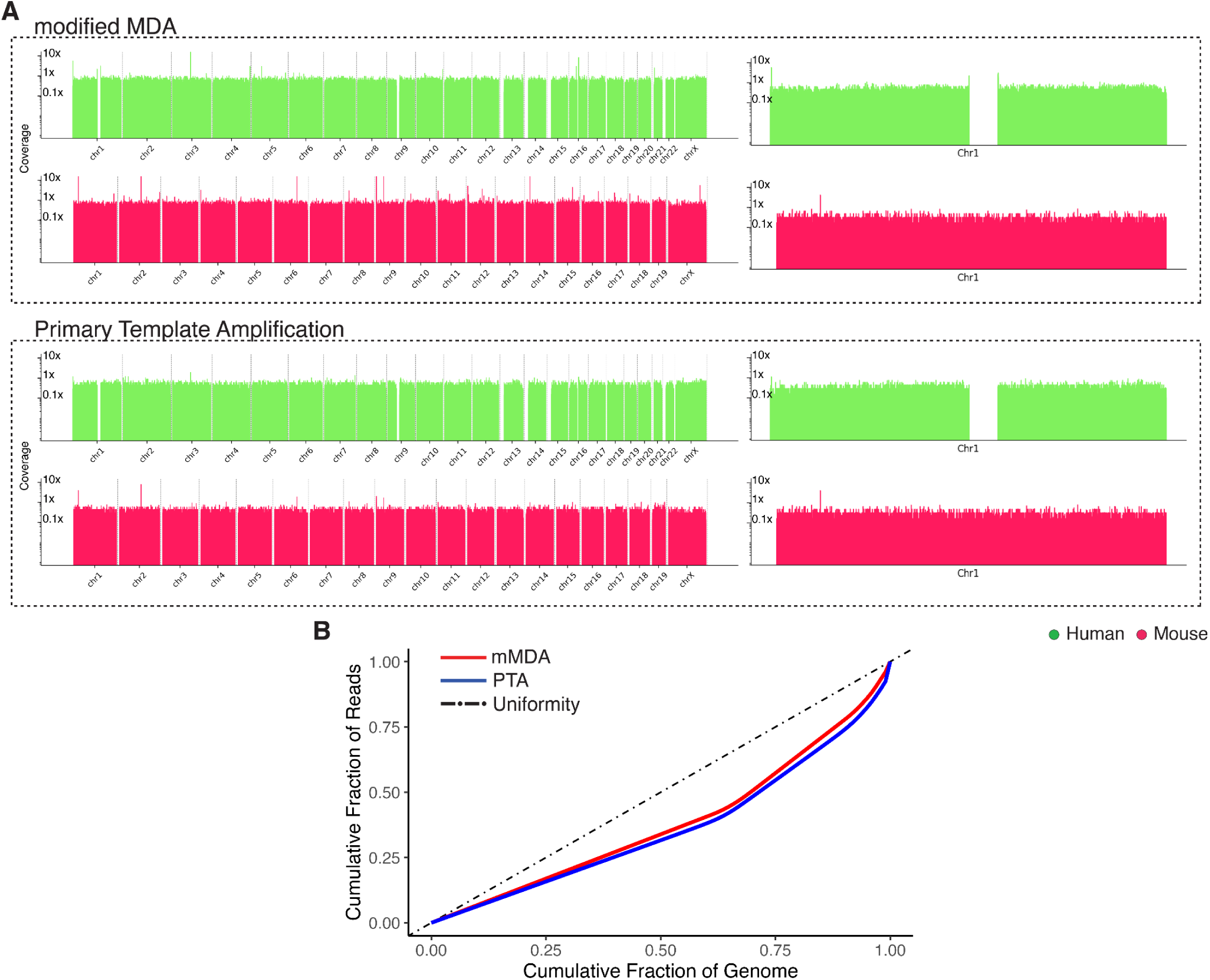
Comparison of whole genome amplification methods. (**A**) Two cells were randomly selected for display from mMDA amplified genomes (top) or PTA amplified genomes (bottom). Whole genome coverage is shown on the left, with an inset of chromosome 1 right. Plots denote coverage within the genome binned at 1kB resolution. Green pile ups denote human cells while red pile ups denote mouse. (**B**) Lorenz curves for modified MDA (mMDA) and primary template amplification (PTA). Dot-dashed line included to indicate optimality or uniformly distributed reads.

### Lineage tracing using the POLE P286R hypermutator allele

The prevalence of somatic evolution — the ongoing mutation and selective expansion of genomic variants within the somatic cells of multicellular organisms — is likely significantly underestimated due to the inherent limitations of bulk WGS approaches (*22*). SPC-enabled sc-WGS can address these limitations by blending uniform coverage with throughput. Nonetheless, reliably resolving naturally arising somatic mutations at single-cell resolution remains challenging, primarily because these mutations are rare, occurring at a rate between 1.6 x 10^-8^ to 7.9 x 10^-10^ mutations per base pair per mitotic division depending on the tissue (*23*). Single base substitutions (SBS), the most common mutation class, reflect intricate interplay between intrinsic cellular processes—such as DNA replication fidelity and repair—and extrinsic environmental exposures that damage DNA. Certain SBS mutational signatures accumulate with chronological aging (e.g., SBS5), whereas others correlate directly with cell division rates or tissue-specific replication processes (e.g., SBS1) (*24*). However, precisely resolving these signatures in single cells remains challenging due to their low frequency and the confounding effects of cell type and cellular differentiation on mutation rate (*25*).

One requirement for genomic lineage tracing — the reconstruction of cellular genealogies based upon heritable genomic alterations — is the consistent, measurable deposition of mutations during successive cellular generations. Historically, naturally occurring somatic mutations have been leveraged to estimate lineage relationships, including seminal work in tracing cancer phylogenies through WGS of clonal somatic variants (*26*). More recently, inducible mutagenesis-based lineage tracing methods, such as those employing CRISPR-Cas9 barcode editing (*27*, *28*) or barcode writing via Prime Editing (*29*) have provided rich, scalable lineage data, however, guaranteeing a regular cadence of barcode deposition has remained difficult.

At the core of eukaryotic genome replication are two major DNA polymerases, polymerase ε (Pol ε) and polymerase δ (Pol δ), responsible respectively for leading and lagging strand synthesis (*30*). Disruption of the proofreading exonuclease domain of these polymerases, particularly via specific amino acid substitutions, can elevate the genome-wide mutation rate by orders of magnitude (*31–33*). Mutations in the Pol ε exonuclease domain, such as POLE P286R, were originally characterized in hypermutated endometrial and colorectal cancers (*26*, *31*), where they yield extraordinarily high mutation burdens exceeding 100 mutations per megabase—the highest observed in human tumors (*34*). Critically for lineage tree time calibration, the mutations generated by proofreading-deficient replicative polymerases usually occur synchronously with DNA replication, thus acting as an intrinsic “molecular clock” that can encode the ages of cellular ancestors.

To test the suitability of POLE P286R for continuous time-calibrated lineage tracing using SPC-based sc-WGS, we introduced the POLE P286R mutation into K562 cells (K562 POLE^P286R^) using CRISPR-Cas9-mediated homology-directed repair. Editing rates exceeded 50% within three days post-editing, as quantified by amplicon sequencing of the targeted locus (**Fig. S14**). Edited cells exhibited no observable fitness disadvantage, maintaining consistent growth rates without apparent selective pressures against the POLE P286R allele over 12 days in culture (**Fig. S14**). Consistent with expectations from previous studies, whole genome sequencing at regular intervals demonstrated a characteristic spike in the TCT>TTT mutation signature and a constant accumulation of mutations, occurring at a rate of 862 mutations per million reads per day (Pearson’s r 0.58), which was significantly elevated relative to controls (**Figure 4B,C**).

**Figure 4.**
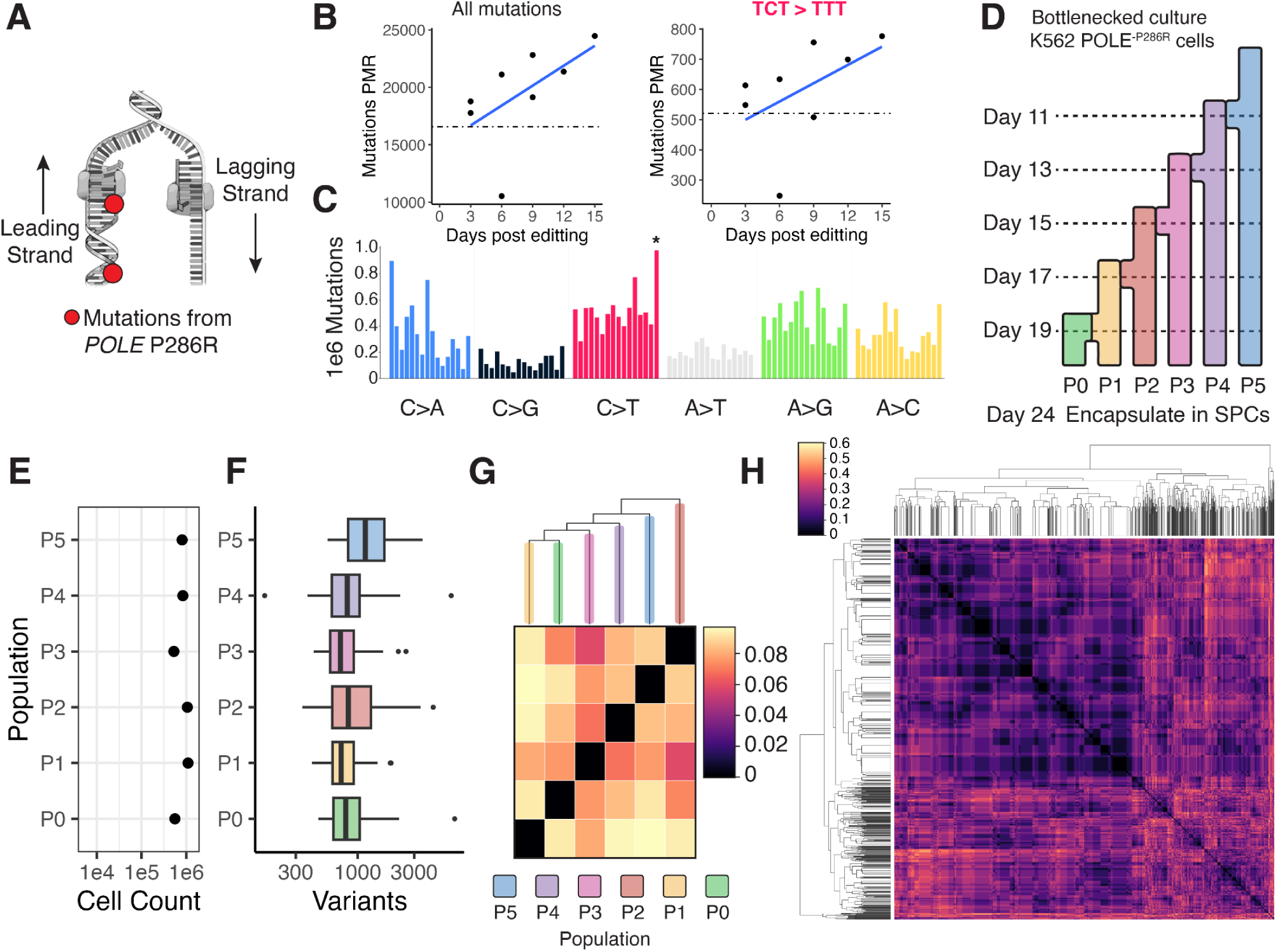
POLE P286R is a continuous recorder of cellular division and cellular lineage. (**A**) Model of the mutations generated by POLE P286R on the leading strand. (**B**) Mutations in genomic DNA per million reads (PMR) are plotted as a function of days in culture for all mutations (left) or TCT>TTT (right). Dotted line is the average number of mutations called in the non-targeting control. (**C**) Trinucleotide context of the mutations detected in POLE P286R after subtraction from mutations detected in the control. POLE P286R trinucleotide signature noted with a (*). (**D**) A small bottlenecked well of K562 POLE^P286R^ cells was split on days as indicated by the diagram and collected 24 days after editing. (**E**) Endpoint cell count of each serially bottlenecked K562 POLE^P286R^ population. (**F**) Variants (with Allele Depth ≥ 2) called per cell plotted as a function of the by population. (**G**) Dendrogram computed by grouping variants within each population and comparing the normalized hamming distances between all sites observed in both groups. (**H**) Single cell lineage structure of 1000 K562 POLE^P286R^ cells, computed by comparing the normalized hamming distance of sites common to pairings of individual cells. Colors in the heatmaps denote normalized hamming distances.

**Supplementary Figure 14.**
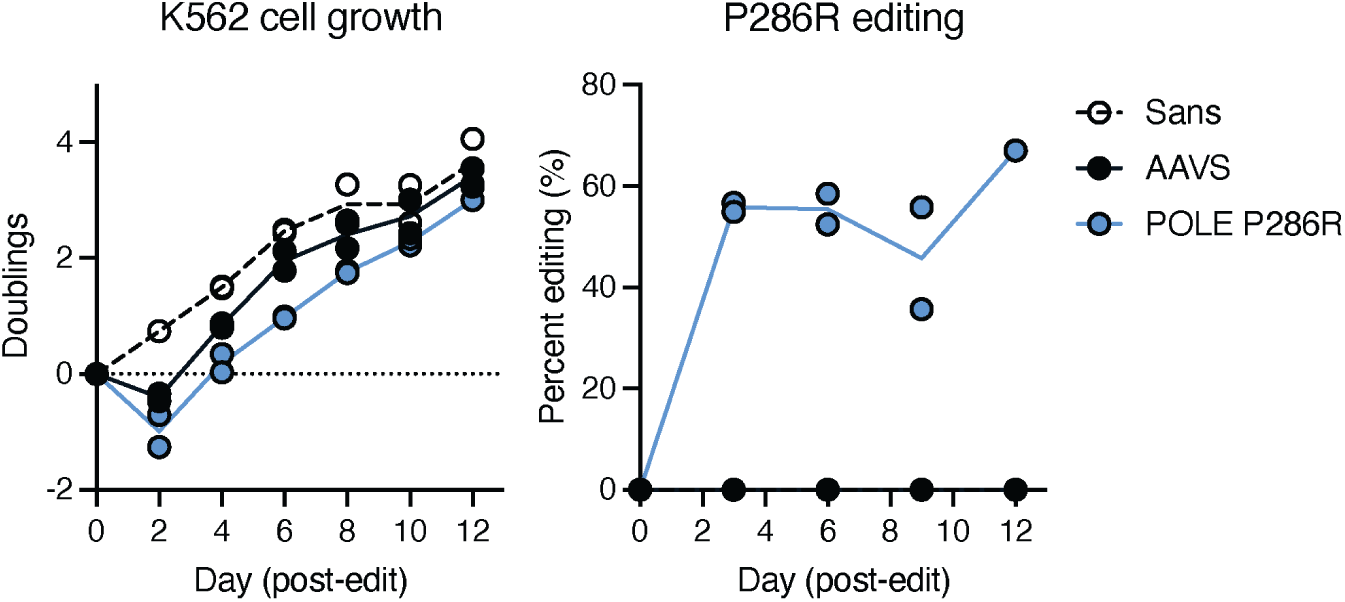
Growth and POLE P286R editing rates of K562 cells. K562 cells were nucleofected with sgRNA targeting POLE and donor template encoding the POLE P286R mutation. Cells were monitored for growth rate (left) and the percentage of templates that were edited (right) versus no editing (Sans) or while using a control sgRNA targeting the safe harbor locus AAVS1.

Given the predictability of mutational accrual associated with the POLE P286R allele, we reasoned that this genetic background could serve as a robust method for high-resolution lineage tracing. To test this concept empirically, we initiated a lineage tracing experiment by subjecting K562 POLE^P286R^ cells to serial bottlenecking, beginning with small founding populations (starting with approximately 3–5 cells), and subsequently expanding and serially splitting these populations to impose a defined lineage structure (**Figure 4D**). Populations proliferated for equal durations (24 days), consistently reaching a density-dependent growth plateau indicative of nutrient and spatial limitation (**Figure 4E**). SPC-based sc-WGS (**Fig. S15**) with shallow sequencing yielded a total of 1,379,244 detectable variants with 2 or more reads supporting the alternative allele, and an average of 954 variants per cell (**Figure 4F**). With the exception of population P2, phylogenetic reconstruction based on these mutational patterns faithfully recapitulated the ground-truth lineage structure (**Figure 4G,H**), demonstrating the POLE P286R allele’s utility for quantitative lineage tracing. The lack of concordance to the anticipated lineage structure by population P2 is possibly attributable to incidental mutations in cell-cycle regulatory genes, and subsequent accumulation of a small subclonal lineage and warrants further investigation.

**Supplementary Figure 15.**
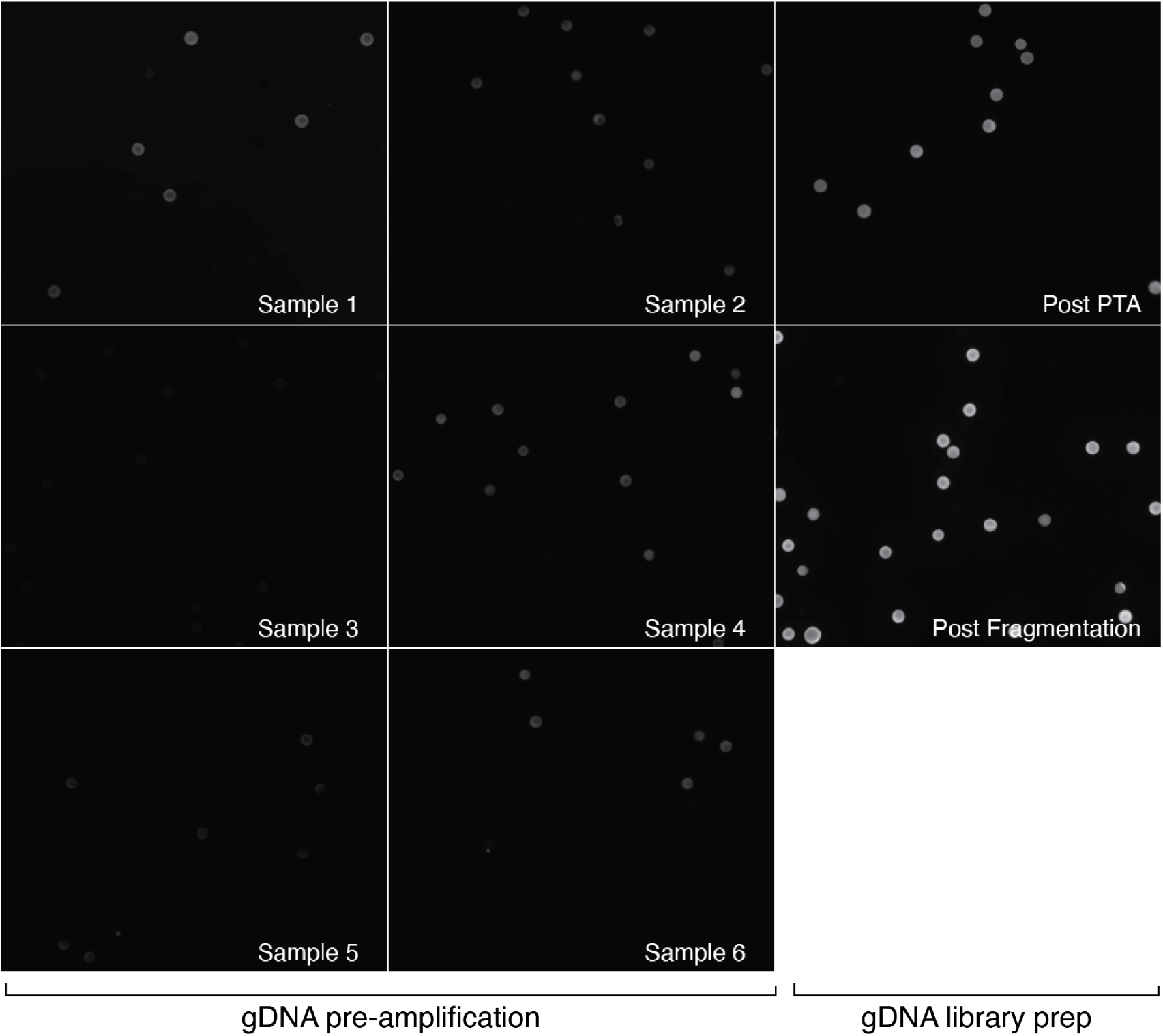
DNA content during library prep. SPCs were stained with SYBR green and imaged at various points during the library prep procedure including after lysis and before amplification (1st and 2nd column), after amplification (third column, top row), or after fragmentation and combinatorial indexing (third column, middle row).

## Discussion

Life’s compartmentalization of matter fundamentally enables the chemical reactions necessary for complex cellular processes and organismal forms (*35*). At the molecular scale, these reactions are governed by information encoded in the structure and sequence of biopolymers, whose identities, arrangements, and stoichiometries have been propagated through successive cell divisions. Against this backdrop, the genome acts as a blueprint for reproducing and replacing molecular machinery, and innovating new biomolecules through mutational processes (*36*, *37*). Inadvertently, these processes leave scars in the genome creating a history of mutational events over time (*38*). Deciphering these genomically-encoded histories within individual cells pinpoints the mechanisms underlying the evolution of both molecules and organisms, providing insight into both form and function.

The study of compartmentalized biological systems requires a compartment-centric experimental approach. In this study, we introduce and characterize a novel commercially available semi-permeable capsule (SPC) comprising a polysaccharide-based shell polymer. These hydrogel capsules can be imagined as picoliter-scale reaction chambers that are capable of containing the contents of a cell for measurement, manipulation, and analysis *ex situ* (*9*). SPCs can withstand a battery of organic solvents, extreme temperatures, and are stable upon treatment with various nucleases and proteases. These properties, in conjunction with the diffusion of biomolecules under a specific size-cutoff, make SPCs a generalizable container for molecular biology, cellular biology, and genomics, enabling a diverse set of applications. We demonstrate the utility of capsule-based assays by growing over 500,000 bacterial or mammalian cultures simultaneously within SPCs, and generating thousands of single cell whole genome sequencing libraries in a single experiment.

Our results illustrate the unique capability of SPCs to maintain high specificity and sensitivity during the amplification of nucleic acids originating from distinct cells. For instance, in single-cell amplicon sequencing assays utilizing multiplexed PCR primers, we successfully detect reads from target amplicons in over 95% of cells, with more than 84% of cells demonstrating the detection of both alleles at a known heterozygous site. Additionally, single-cell whole-genome amplification and sequencing within SPCs yielded libraries with highly uniform genome-wide coverage and minimal cross-contamination between species. We then leveraged this capability by conducting large-scale single cell whole-genome sequencing of 1,000 bottlenecked K562 cells harboring a hypermutator allele of polymerase ε (POLE P286R), the enzyme responsible for leading-strand DNA synthesis. The constant accumulation of mutations genome-wide in these cells when deployed in conjunction with single cell whole genome sequencing, can be used to reconstruct cellular lineages and estimate the number of cell divisions a cell has undergone.

We anticipate that the use of molecular recorders linked to constant processes like cell division, will enable the investigation of a new class of biological questions that have been previously inaccessible in optically opaque organisms. Such questions range from the extent of dominance exhibited by cellular clones during physiologic processes such as the formation of organs, tissue regeneration, or disease (*39*). Intriguingly, mice carrying the hypermutator and ultramutator alleles are viable, undergoing normal development despite their high somatic mutation rates and thus providing a path towards performing lineage tracing *in vivo* . The access to both flexible methods for generating single cell genome libraries at scale, in conjunction with the continued scaling of next generation sequencing technologies (*40*, *41*) promise to make these fundamental questions ones that will be addressable in the near future.

## Acknowledgements

We thank members of the Srivatsan lab for critical reading and feedback. This research was supported by the Genomics & Bioinformatics Shared Resource (RRID:SCR_022606), Cellular Imaging Shared Resource (RRID:SCR_022609) and Electron Microscopy Shared Resource (RRID:SCR_022611) which are funded by the Fred Hutch/University of Washington/Seattle Children’s Cancer Consortium (P30 CA015704). Part of this work was also conducted at the Molecular Analysis Facility, a National Nanotechnology Coordinated Infrastructure (NNCI) site at the University of Washington, which is supported in part by funds from the National Science Foundation (awards NNCI-2025489, NNCI-1542101), the Molecular Engineering & Sciences Institute, and the Clean Energy Institute. Assistance for computing provided by FHCC IT - Scientific Computing team, supported by NIH grants S10-OD-020069 and S10-OD-028685.

## Author Contributions

S.R.S., K.H., and R.Ž. were involved in the initial planning and conceptualization of the project. S.R.S., S.R.S., D.M., I.C., L.K., and A.Š. performed the majority of the experiments. D.B.M., and Y.H., conducted the analyses and designed the genomics analyses with input from S.R.S and K.H. Both H.E, and J.S. designed and cloned *de novo* designed proteins. V.E.B., E.K., S.S. and S.R.S. performed image processing and image analysis. S.R.S., D.B.M, S.R.S, and D.M. wrote the manuscript with input from all co-authors.

## Data and materials availability

All data and code will be uploaded to public repositories and this preprint will be updated shortly to include links to both.

## Supplementary Files

**Supplementary Video 1. Video of 65µm SPC generation.** SPCs generation visualized on the Onyx Droplet Microfluidic platform (Atrandi Biosciences). Video captured at a frame rate of 50µs and rendered at 30 frames per second.

**Supplementary Video 2. Capsule Release After Addition of Glycosidase .** HEK293T cells encapsulated in semi-permeable capsules. 10μL of capsules were released via addition of 1μL of Release Agent, at t=0. Capsules were imaged at 20× magnification on a Nikon Live imager; scale bar:50µm

## Methods

### Generation of SPCs using Co-Flow Devices

A complete description of flow rates, microfluidic chip characteristic sizes, and product numbers, can be found in **Table S1** for the generation of capsules at a range of sizes. Below we have outlined a description for the generation of 200µL of 65µm ± 5µm capsule suspension. Volumes of Core Solution (CS), Shell Solution (SS), and Capsule Stabilization Oil (CSO) can be scaled based on flow rates for each device or the batch size of capsules desired.

SPCs were generated by making CS and SS separately prior to loading the solutions and CSO into the microfluidic chip. To make CS, 50µL of Core (CRP-CR1: Atrandi Biosciences), 36.5µL of 1x PBS (70011-044: Gibco), 12.5µL of Photoinitator (CRP-PA1: Atrandi Biosciences), and 1µL of 1M DTT (646563: Sigma-Aldrich) were combined in a microcentrifuge tube. <u>This formulation is referred to as the standard core solution</u>, general protocols replace the 36.5 PBSµL of 1x PBS with an experiment appropriate buffer or material for encapsulation. To make SS 50µL of Shell (CRP-SR1: Atrandi Biosciences) and 50µL of 1x PBS (70011-044: Gibco) were combined in a microcentrifuge tube. <u>This formulation is referred to as the standard shel solution.</u> Both CS and SS were mixed by gentle pipetting, attempting not to introduce bubbles followed by centrifuging for 1 minute at 13,000*g on a fixed angle benchtop centrifuge. For consistency, Core and Shell solutions were pipetted and mixed using wide bore tips or pipette tips cut with a razor blade.

Both SS and CS were loaded separately into 1mL syringes with a 400µL underlay of Sample Loading Oil (MON-SLO1: Atrandi Biosciences). Finally, 750µL of CSO was loaded into a 1mL syringe. All three syringes were primed and connected to a co-flow microfluidic chip. Emulsions were generated after setting flow-rates (specific values in **Table S1**) and collected in a 1.5mL microcentrifuge tube. After collection, emulsions were inverted gently five times and then polymerized for 30 seconds using 405nm light (MHT-LAS1: Atrandi Biosciences). After polymerization, emulsions were broken with 500µL of a 1:1 mixture of Emulsion Breaker (MON-EB1: Atrandi Biosciences) and 1x PBS, containing 1x Wash Additive (CRP-WA1: Atrandi Biosciences). The aqueous layer (top layer) containing the SPCs, was then moved to a new 1.5mL microcentrifuge tube and washed once again with 500µL of emulsion breaking solution. SPCs were stored in 1x PBS augmented with 0.1x Triton-X (X100-100ML: Sigma-Aldrich). Triton-X prevents the SPCs from sticking to the walls of the plastic tube.

**Supplementary Table 1.**

| Chip | Expected Size | CS flow rate<br>(μL/hour) | SS flow rate<br>(μL/hour) | CSO flow rate<br>(μL/hour) |
| --- | --- | --- | --- | --- |
| MCN-C2 | 35μm | 25 | 25 | 300 |
| MCN-C4 | 65μm | 75 | 75 | 450 |
| CCN-G4-Q2 | 175μm | 150 | 150 | 600 |

### Chemical Stability Experiments (Figure S1)

SPCs were loaded with 2 million HEK-293T cells / mL, polymerized, and washed before lysis with GeneJet cell buffer (Thermo Fisher; K0731). Genomic DNA was amplified using primers (247F, CCGCCGCTTTCCTTAACCACAAATCAGGC) and (255R, TGCACCACTTCCCAGGACATAGGCGTGTG).

Targets were amplified using the following program: 1 cycle of 98°C for 3 minutes; 40 cycles of (98°C for 20 seconds, 65°C for 20 seconds, 72°C for 1.5 minutes); and a final extension at 72°C for 5 minutes, using Kapa HiFi Hotstart Readymix (Roche). After PCR, SPCs were prepared for either overnight organic solvent, acid/base, or temperature exposure. In the organic solvent condition, SPCs were solvent exchanged by washing twice in 100% ethanol and were charged into a 1-dram glass vials (Fisher) and incubated in hexanes (Sigma Aldrich), diethyl ether (Fisher), dichloromethane (EMD Millipore), 2-propanol (Sigma Aldrich), 1-propanol (Sigma Aldrich), ethanol (Sigma Aldrich), ethyl acetate (Fisher), acetone (Fisher), methanol (Fisher), N-N,dimethylformamide (Fisher), or dimethyl sulfoxide (Fisher). SPCs were recovered from overnight incubation into 100% ethanol and washed twice with 100% ethanol. In the acid/base condition, SPCs were washed twice in ddH_2_O and were charged into a 1-dram glass vial and incubated in dilutions of NaOH or HCl. In the temperature conditions, SPCs were incubated with 1× Wash Buffer (CRP-WA; Atrandi Biosciences) during all temperature treatments. In all temperature conditions (-196°C, -80°C, 22°C, and 100°C) SPCs were incubated for 1 hour. In the -196°C condition, a microcentrifuge tube containing SPCs was placed on a float rack and allowed to sit on liquid nitrogen. In the -80°C condition, a microcentrifuge tube containing SPCs was placed in a cryogenic freezer. Room temperature SPCs were allowed to sit benchtop. Finally, in the 100°C condition, a microcentrifuge tube with SPCs was placed on a preheated, benchtop heat block. Before imaging, SPCs were washed twice with hypotonic nuclei buffer (3mM MgCl_2_, 10mM Tris-HCl, 10mM NaCl, 0.1% Tween-20, and 0.1% NP-40) and stained with SYBR Gold.

### Physical SPC Disruption

SPCs were broken using high impact 1.5mm zirconium beads (Benchmark Scientific; D1132-15TP). First, 5-20 zirconium beads were added to 20-150µL of SPCs, in a 1.5mL microcentrifuge tube. SPCs and beads were then vortexed at maximum speed for 1 minute. SPCs and beads were then passed over a 5µM cell strainer (PuriSelect; 43-10005-40), at 1,000*g for 1 minute, into a fresh 1.5mL microcentrifuge tube.

### Microscopic Imaging of SPCs

For most experiments, SPCs were visualized in Countess cell counting chamber slides (C10228: Thermo Scientific). For larger SPCs that did not fit within cell counting chamber slides, SPCs were placed on a cover glass bottom 24 well plate (CellVis; P24-1.5H-N) prior to imaging. Most images were acquired on an Echo Revolve microscope with the following objectives:

| Objective | NA | Pixel Size (µm/pixel) | Phase Ring |
| --- | --- | --- | --- |
| 4x | 0.13 | 0.8838 | No Phase |
| 10x | 0.30 | 0.3535 | Phase 1 |
| 20x | 0.45 | 0.1768 | Phase 1 |

### SEM (Figure 1C, Figure S2)

Cells/coverslips/HMDS for SEM. Duplicates of 50µLof each sample were applied in a pool on poly-l-lysine coated coverslips for 30min and then rinsed twice with 0.1M sodium cacodylate buffer. The coverslips were then dehydrated through a graded series of alcohols, infiltrated with several changes of HMDS (Electron Microscopy Sciences, Hatfield, PA) and allowed to air dry. Coverslips were then mounted on aluminum stubs and sputter coated with gold/palladium (Denton Desk IV, Denton Vacuum, Moorestown, NJ). Samples were imaged on a JSM 6610 LV scanning electron microscope at 15kV (JEOL, Tokyo, Japan).

### Live Imaging and Capsule Release (Supplemental Video 2)

10µL of either empty, encapsulated K562, or encapsulated HEK293T SPCs in 40µL of 1 x PBS were loaded onto a 0.17mm (+/- 0.005 mm) glass bottom plate, and allowed to settle. Videos were taken on a Nikon Live imager. To release capsules, 1µL of Release Agent (Atrandi; CRP-PA1) was added to the end of a capillary tube. Capillary was placed at the center of each well, and at t = 0, 1 x PBS was gently injected at the opposite end of tubing, to introduce Release Agent into the well. Videos were captured using bright field (BF) and GFP channels, using 20X objective.

### Retention Experiments (general) (Figure 1E,F, Figure S3)

SPCs were generated with the MCN-C4 chip. After SPC formation, emulsions were broken by the addition of 200µL of 1H,1H,2H,2H-Perfluorooctanol (ThermoFisher; B20156.18) followed by quick vortexing. Broken emulsion was then spun at 1000g for 30 seconds, and residual oil underneath the SPC layer was removed. To rehydrate SPCs, 150µL of 1x PBS was added to the solution and the solution was quickly vortexed. SPCs were spun down and top, aqueous layer was captured. 40µL of SPCs were then collected as the t = 0 fraction. Next, 850µL of 1x PBS was added to SPCs, and the mixture was allowed to incubate with end-over-end rotation until the next time point. At each time point, SPCs were centrifuged at 2000*g for 1 minute, supernatant was discarded, and a 100µL aliquot of hard packed SPCs was recovered. This procedure was repeated and samples were collected at the 5 minute time point (1 wash), 10 minute time point (2 washes), 30 minute time point (3 washes) and 60 minute time point (4 washes). After the final collection, approximately 5 1.5mm Zirconium beads (Benchmark Scientific; D1132-15TP) were added to each sample and vortexed for 30 seconds to physically disrupt the SPCs. Fragmented SPCs and beads were then passed over a 5µM cell strainer (PuriSelect; 43-10005-40), at 1,000*g for 1 minute, into a fresh 1.5mL microcentrifuge tube. Finally, purified flowthrough was run on a gel depending on the sample type.

### DNA Ladder Retention (Figure 1E)

Core: 25µL of TrackIt 1kB Plus DNA ladder (Invitrogen; 10488085) and 12.5µL of Ultra Low Range DNA Ladder (Thermo Fisher; 10597012) were mixed into the standard core working solution. Shell working solution was standard composition and 65µm SPCs were generated using standard flow rates, with the MCN-C4 chip. After physical disruption of SPCs, 15µL of product was mixed with 4µL of 6x loading dye and run on a 6% TBE gel. After completion, the gel was stained with SYBR gold and imaged on the Biorad Chemidoc.

### Protein Ladder Retention (Figure 1F, Figure S3)

Native Protein Standard Mix 15 - 600 kDa (Millipore Sigma; 69385-30MG) was reconstituted in 1mL of water.

36.5μL of this protein solution was loaded into the core solution. The shell solution was standard composition. 65µm. SPCs were generated with standard flow rates, with the MCN-C4 chip. After physical disruption of the SPCs, 100µL of 2x Laemmli Sample Buffer (Biorad; #1610737), supplemented with 2-Mercaptoethanol, was added to each sample and boiled for 5 minutes at 80°C. 15µL of each sample was run on a Mini-PROTEAN TGX Stain-Free gel (Biorad; #4568126), stained with coomassie Blue, and imaged.

### Mononucleosome Purification and Retention Experiment (Figure 1F, Figure S3)

Nucleosomes were isolated from K562 cells. K562 cells were crosslinked in ice-cold crosslink buffer (80mM PIPES-KOH (pH 6.8), 15mM NaCl, 60mM KCl, 30% Glycerol, 1mM MgCl_2_, 10mM β-glycerophosphate, 10mM Sodium Butyrate, 1% formaldehyde). Crosslinked cells were permeabilized by digitonin buffer (80mM HEPES-KOH (pH 7.4), 15mM NaCl, 60mM KCl, 0.05% Digitonin, 50mM Glycine, 1.25x Protease inhibitor, 10mM β-glycerophosphate, 10mM Sodium Butyrate, 1mM MgCl_2_, and 5mM CaCl_2_) and treated with 0.9µM MNase at 4°C for 16 hours. The mononucleosome-containing fraction was isolated by 12-22% sucrose gradient (*42*) and 36.5µL of the mononucleosome fraction was mixed into the CS. After collection of fractions from the retention testing, mononucleosomes were either treated with 0.5µL of Thermolabile Proteinase-K (P8111S: New England Biolabs) for 30 minutes at 37°C or not. 10µL aliquots of these samples were then run on a native 6% TBE gel and visualized after staining with SYBR Gold.

### Ribosome Retention (Figure 1F, Figure S3)

Ribosomal subunits were purified from HEK293T as described in ref (*43*). Ribosomes were reconstituted by mixing purified 40S and 60S ribosomal subunits in 80S formation buffer [150mM KOAc, 200mM MG(OAC)^2^, 30mM HEPES-KOH (pH 7.4), 400mM Sucrose and 200mM DTT] for 10 minutes at 37 C to form the 80S particle. 36.5μL of reconstituted ribosome was then loaded into the core. The shell solution was standard composition. 65µm SPCs were generated with standard flow rates. After physical disruption of the SPCs, 100µL of 2x Laemmli Sample Buffer (Biorad; #1610737) supplemented with 2-Mercaptoethanol was added to each sample and boiled for 5 minutes at 80°C. 15µL of sample was run on a Mini-PROTEAN TGX Stain-Free gel (Biorad; #4568126), stained with coomassie stain solution, and imaged.

### Protein Design (Figure 1F, Figure S3)

Tetrahedral and icosahedron nanoparticle backbones of 80 amino acids per subunit were generated with RoseTTAFold Diffusion as previously described (*11*). Following diffusion, backbones were minimally downsampled, only filtering on external C-termini (for experimental purification) and inter-subunit contacts to ensure sufficient interface size. Backbones were then designed as homo-oligomers with ProteinMPNN at a sampling temperature of 0.1 and with 8 sequences per RFdiffusion-generated backbone (*44*). Candidate sequences were then predicted (asymmetric unit only) with AlphaFold2 (*45*) and designs were filtered on pLDDT ≥ 85 and RMSD ≤ 1 Å, with 1,187 designs passing.

### Protein Purification (Figure 1F, Figure S3)

Glycerol stocks (100µL) were used to inoculate 50mL Terrific Broth in 250 mL baffled flasks. Cultures were grown for 8h at 37°C before the addition of IPTG to a final concentration of 0.1mM. Bacteria were then grown overnight at 37°C and cell pellets were harvested via centrifugation (30 min, 4000 x g). Pellets were resuspended in 25 mL lysis buffer (25 mM Tris HCl pH 8, 300 mM NaCl, 40 mM Imidazole, 1 mM DNase I, 10 µg /mL lysozyme) and lysed by sonication (Q500 Sonicator Dual Horn ¾ ” probes, Qsonica, 5 min, 85% amplitude, 15s on/off cycles). Lysate was clarified by centrifugation (14,000 x g, 20 min.). The supernatant was run over the 5mL of Ni-NTA Agarose (Qiagen; 30230). The column was then washed with 2 column volumes of Wash buffer (25 mM Tris HCl pH 8, 300 mM NaCl, 40 mM imidazole) and then eluted in 3mL of elution buffer (25 mM Tris HCl pH 8, 150 mM NaCl, 400 mM imidazole). Proteins were then dialyzed into 1x PBS before performing size exclusion chromatography (SEC). SEC was performed on an equilibrated Superdex 200 Increase 10/300 GL at a flow rate of 0.3 mL/min in1x PBS. Fractions were collected in 0.25 mL intervals from 4 mL to 40 mL. The corresponding fractions for the oligomer were combined and then concentrated using a centrifugal concentrator with a 3kD molecular weight cutoff (Millipore Sigma; UFC900308).

### Protein Design Retention (Figure 1F, Figure S3)

36.5µL of each concentrated protein design was loaded into the core solution. The shell solution was standard composition. 65µm SPCs were generated with standard flow rates. After physical disruption of the SPCs, 100µL of 2x Laemmli Sample Buffer (Biorad; #1610737) supplemented with 2-Mercaptoethanol was added to each sample and boiled for 5 minutes at 80°C. 15µL of sample was run on a Mini-PROTEAN TGX Stain-Free gel (Biorad; #4568126), stained with coomassie Blue, and imaged.

### Nalm6 Staining in SPCs (Figure 1G)

For staining experiments, Nalm6 cells were encapsulated in SPCs at 1.5 × 10⁶ cells per 100µL of SS as described above. SPCs were incubated at room temperature with minimal light exposure at a 1:100 dilution of anti-CD19-PE (Biolegend) for a specified time. Cells were then pelleted by centrifugation at 500 × *g* for 3 minutes, and washed twice with FACS buffer (PBS supplemented with 10% FBS and 1mM EDTA). Final solutions of 200µL SPC-Nalm6 were released by the addition of 4µL of release agent. Cells were passed through a 22µM filter and fluorescence was then quantified via flow cytometry on the Cytek Aurora.

### Bacterial Culture (Figure 2B-D)

SPCs encapsulating bacterial cells were generated as follows. To prepare the **core solution (CS)**, the following components were combined in a microcentrifuge tube: 50µL of Core reagent (CRP-CR1; Atrandi Biosciences), up to 36.5µL of Terrific Broth (TB) (Thermo Fisher; A1374301), log phase *E. Coli* in TB, 12.5µL of Photoinitiator (CRP-PA1; Atrandi Biosciences), and 1µL of 1 M DTT (64656; Sigma-Aldrich). For the **shell solution (SS)**, 50µL of Shell reagent (CRP-SR1; Atrandi Biosciences) was mixed with 50µL of TB. When targeting sub-Poissonian loading of single cells per capsule, SPCs were generated with a total of 314,000 cells suspended in 50µL of conditioned media, yielding an estimated recovery of ∼200,000 cells and estimated multiplet rate in SPCs of 11.3%. Immediately after encapsulation, 500µL of emulsion breaker solution (MON-EB1; Atrandi Biosciences) was added to the mixture along with 500µL of TB. The mixture was inverted 3 times and incubated for 3 minutes at room temperature, followed by centrifugation at 300 × *g* for 3 minutes to separate the loading oil from the SPCs. The oil layer was removed via pipetting and discarded. SPCs were then grown in 1.5mL of TB in a 14-mL round bottom tube for 8 hours before imaging on a Countess slide. For bacterial growth experiments, 10μL of SPCs were broken through the addition of 0.5µL of Release Agent (Atrandi; CRP-PA1) and plated on agar at multiple dilutions. After growth colonies were counted and used to estimate the growth rate.

### Cell Culture (Figure 2E-M, Figure S6-S9)

Cells grown in suspension, such as K562, Nalm6, and Jurkat cells, were cultured in RPMI (Gibco) supplemented with 10% FBS (Cytiva), 1% penicillin and streptomycin (Gibco), and 1% GlutaMAX (Gibco). Adherent cells, such as HEK293T cells, were cultured in DMEM (Gibco) supplemented with 10% FBS, 1% penicillin and streptomycin, 1% HEPES (Gibco), and 1% GlutaMAX.

### Generation of SPCs Encapsulating Mammalian Cells (Figure 2E-M, Figure S6-S9)

SPCs encapsulating mammalian cells were generated by adapting previously established parameters. All SPCs were prepared by separately formulating the core solution (CS), shell solution (SS), and capsule stabilization oil (CSO). To prepare the **core solution (CS)**, the following components were combined in a microcentrifuge tube: 50µL of Core reagent (CRP-CR1; Atrandi Biosciences), 28.5µL of 1× conditioned media, 12.5µL of Photoinitiator (CRP-PA1; Atrandi Biosciences), and 9µL of 1 M DTT (64656; Sigma-Aldrich). For the **shell solution (SS)**, 50µL of Shell reagent (CRP-SR1; Atrandi Biosciences) was mixed with 50µL of the suspended cell mixture in a separate microcentrifuge tube. When targeting sub-Poissonian loading of single cells per capsule, SPCs were generated with a total of 750,000 cells suspended in 50µL of conditioned media, yielding an estimated recovery of ∼500,000 cells and estimated multiplet rate in SPCs of 12%. For standard cell growth experiments, cell suspensions containing 1.5–3 × 10⁶ cells in 50µL were used, resulting in an estimated recovery of 1–2 × 10⁶ cells post-encapsulation and an estimated multiplet rate of at least 33%. Immediately after encapsulation, 500µL of emulsion breaker solution (MON-EB1; Atrandi Biosciences) was added to the mixture along with 500µL of conditioned media. The mixture was incubated for 3 minutes at room temperature, followed by centrifugation at 300 × *g* for 3 minutes to separate the loading oil from the SPCs. The oil layer was decanted from the bottom and discarded. To further purify the SPCs, 500µL of additional media supplemented with 1× Wash Buffer (CRP-WA; Atrandi Biosciences) was added. After centrifugation at 300 × *g* for 1 minute, any residual oil was decanted, and the wash step was repeated once more. Cell-containing SPCs were then plated with 1:1 conditioned media to fresh media.

### Cell Counting Experiments (Figure 2E-J, Figure S6A-D, S7)

For all cell lines, we generated stable expressing cells to facilitate downstream imaging and counting during SPC culture. Briefly, we produced lentivirus harboring an EGFP expression cassette for integration at the AAVS1 locus. Following transduction, fluorescence was confirmed via flow cytometry and EGFP-positive cells were sorted. After a recovery period, a subset of these cells was used for encapsulation, as described above. To assess cell proliferation, we compared the growth rates of encapsulated cells to those of free-floating cells of the same passage and density, ensuring a similar initial number of cells per cm² of culture surface area. All conditions were plated in triplicate. Growth rates were determined by counting a fixed volume of cells in both SPC and free cultures of vigorously mixed culture was taken from each condition. In the case of SPC cultures, a ratio of 1µL of release agent per 20µL culture was added, followed by a 3-minute incubation at room temperature. A 10µL sample of the released cell suspension or undisturbed cells was then mixed with 10µL of trypan blue to assess viability. Live cells were quantified using a Countess 3 Automated Cell Counter (ThermoFisher Scientific).

### Cell Freezing Experiments (Figure 2L,M)

For freezing experiments, SPCs were generated with a high-density loading of 3 × 10⁶ K562 cells per 100µL of SS. Following encapsulation, cells were allowed to rest for 24 hours before being frozen in one of three cryopreservation formulations: 0%, 5%, or 10% DMSO in fresh RPMI media supplemented with 15% FBS. For each condition, approximately 225,000 SPCs—corresponding to 425,000 cells per vial—were aliquoted in triplicate into vials containing a final volume of 500µL. Cells were gradually cooled to -80°C at a rate of 1°C per minute and stored for 5 days at -80°C. To thaw the cells, vials were quickly brought to room temperature using a 37°C water bath. A total of 500µL of pre-warmed media was added to each vial, and the cells were pelleted by centrifugation at 300 × *g* for 1 minute to remove the freezing media. All conditions were resuspended in 1 mL of pre-warmed media and plated in a 6-well culture-treated plate. For cell viability imaging, cultures were co-incubated with 1× SYTO-9 (at a 1:2000 volumetric ratio) and 1× ethidium homodimer-1 (at a 1:500 volumetric ratio) for 10 minutes at 37°C. Live cells were quantified as described above.

### Fibronectin coating of SPCS (Figure 2K)

For coating experiments, SPCs were assumed to have a 65µm diameter, with a surface area of approximately 13.3 x 10^-4^ cm^2^ per SPC. Fibronectin at 0.05M (F1141; Millipore Sigma) was added to the core mixture at 5 µg / per cm^2^ for approximately 100,000 SPCs per run. A following final composition was used for the CS: 50µL of Core reagent (CRP-CR1; Atrandi Biosciences), 2.95 µL of 1:1000 fibronectin, 22.55 µL of 1× conditioned media, 12.5µL of Photoinitiator (CRP-PA1; Atrandi Biosciences), and 9µL of 1 M DTT (64656; Sigma-Aldrich). For SS, 50µL of Shell reagent (CRP-SR1; Atrandi Biosciences) was mixed with 50µL of a suspended HEK239T cell mixture of containing 1.5 × 10⁶ cells in a separate microcentrifuge tube.

### Confocal imaging for quantification of SPC occupancy by K562 cells (Figure S6 H-I)

SPCs encapsulating fixed K562s were generated by loading cells at a high density and allowing cells to grow in SPCs in culture for 10 days. Following cell growth, SPCs were washed once in 1x PBS before fixing with 4% PFA for 15 minutes at room temperature. Fixative was quenched by adding 1mL of 1M Tris/HCl pH 7.4. SPCs were then washed with 1x PBS. The SPCs were centrifuged at 300g for 3 minutes and resuspended in 6μM Yo-Pro-1 monomeric DNA stain (Biotium Cat. No. 40089) in 1xPBS for 10 minutes. The SPCs were then centrifuged at 300g for 3 minutes and resuspended in 1xPBS. After a final round of centrifugation and rinsing in 1xPBS, 25μL of the stained SPC suspension was placed on a glass coverslip, which was sealed onto a glass slide with nail polish. Confocal imaging of the stained SPCs was performed using Yokogawa CSU-W1 SoRa 15 spinning disc confocal attached to a Nikon Eclipse Ti2 microscope. Excitation light was emitted at 30% of maximal power from a 488nm laser housed by a Nikon LUNF 405/488/561/640NM 1F commercial launch. A single-mode optical fiber transmitted the excitation light to the CSU-W1 SoRa unit. The excitation light was then directed through a microlens array disc and a SoRa disc containing 50µm pinholes and directed the the rear aperture of a 60x N.A. 0.95 Plan Apo lambda air objective by a prism in the base of the Ti2. Emission light was collected by the same objective and passed by the prism back into the SoRa unit, where it was relayed by a 1x lens through the pinhole disc and directed into the emission path by a quad-band dichroic mirror (Semrock Di01- 25 T405/488/568/647-13x15x0.5). Emission light was then spectrally filtered by a bandpass filter (ATTO 488: Chroma ET525/36M) and focused by a 1x relay lens onto an Andor Sona 4.2B-11 camera with a physical pixel size of 11µm, resulting in an effective pixel size of 183.3 nm. The Sona was operated in 16-bit mode with rolling shutter readout and an exposure time of 100ms. Nikon NIS-Elements was used to define a volumetric tiled acquisition (20 FOV x 20 FOV) with 5% tile overlap and 237 z-planes with a z-step of 0.2 um, covering ∼12.7 mm^2 at a depth of 47.4 um for a total volume of ∼ 0.6mm^3. BigStitcher, an ImageJ plugin (https://github.com/PreibischLab/BigStitcher) was used to align the tiles and generate a single stitched image. The ‘cyto3’ model from CellPose 3.0 (https://github.com/MouseLand/cellpose) was used to segment nuclei in the image volume. After generating nuclei segmentation masks, DBSCAN was run to assign nuclei to spatial clusters (SPCs) using the nuclei centroid positions. The clustering results were then manually refined in ImageJ to correct nuclei assignments in sparsely-populated SPCs. Specifically, SPC boundaries were identified by adjusting the image contrast and incorrect cluster assignments were resolved by selecting regions using the ImageJ selection tool and assigning a common label to all nuclei centroids within each region. Finally, the python package *matplotlib* was used to generate a histogram showing the number of nuclei detected within each SPC.

### PCR Amplicon Ladder Experiment (Figure S10)

Overloaded SPCs were generated by creating a solution containing 700,000 cells in 36.5µL 1x PBS. The SPC generation protocol was otherwise standard. Upon generation, SPCs were polymerized, broken and washed before digestion with Proteinase-K in lysis buffer (0.1% SDS, 10mM Tris/HCl pH 7.5). These SPCs were used as template for PCR reactions. PCR reactions were performed with 0.5µM of the forward and reverse primer (**Table S3**) along with Kapa HiFi 2x Master Mix (Roche; KK2602) and 10μL of hard packed SPCs in a 50µLreaction. PCR was performed with the following program:

------------------------------------------------

98°C for 3 minutes

30 cycles of:

> 98°C for 15 seconds
>
> 63°C for 30 seconds
>
> 72°C for 2 minutes

72°C for 5 minutes

------------------------------------------------

Supernatant was recovered by pelleting SPCs and stored. SPCs were washed 5x in 200µL Wash Buffer 1 (WB1) [10mM Tris/HCl pH 7.5, 0.1% Triton-X, 10mM EDTA]. SPCs were stained by adding 1µL of SYBR Green (200x working concentration) to pelleted SPCs with 100µL of 1x PBS. SPCs were imaged with a 4x objective by loading concentrated SPCs onto a countess slide. Gel electrophoresis was performed after breaking 5µL of SPCs by adding 1µL of Release Agent (Atrandi; CRP-PA1). Supernatant and SPC fractions were run on a 6% TBE gel, stained with SYBR Gold and imaged.

### Empty SPC, amplicon swapping experiment (Figure S12)

Empty SPCs were generated by creating a core solution containing 36.5µL 1x PBS. The SPC generation protocol was standard. Upon generation, SPCs were polymerized, broken and washed in 1x PBS. These SPCs were placed in a PCR reaction in addition to 0.5µM of the forward and reverse primer (**Table S2**), Kapa HiFi 2x Master Mix (Roche; KK2602) and genomic DNA at a final concentration of 0.48 ng/µL in a 50μL reaction. PCR was performed with the following program:

------------------------------------------------

98°C for 3 minutes

30 cycles of:

> 98°C for 15 seconds
>
> 63°C for 30 seconds
>
> 72°C for 2 minutes

72°C for 5 minutes

------------------------------------------------

**Supplementary Table 2:**
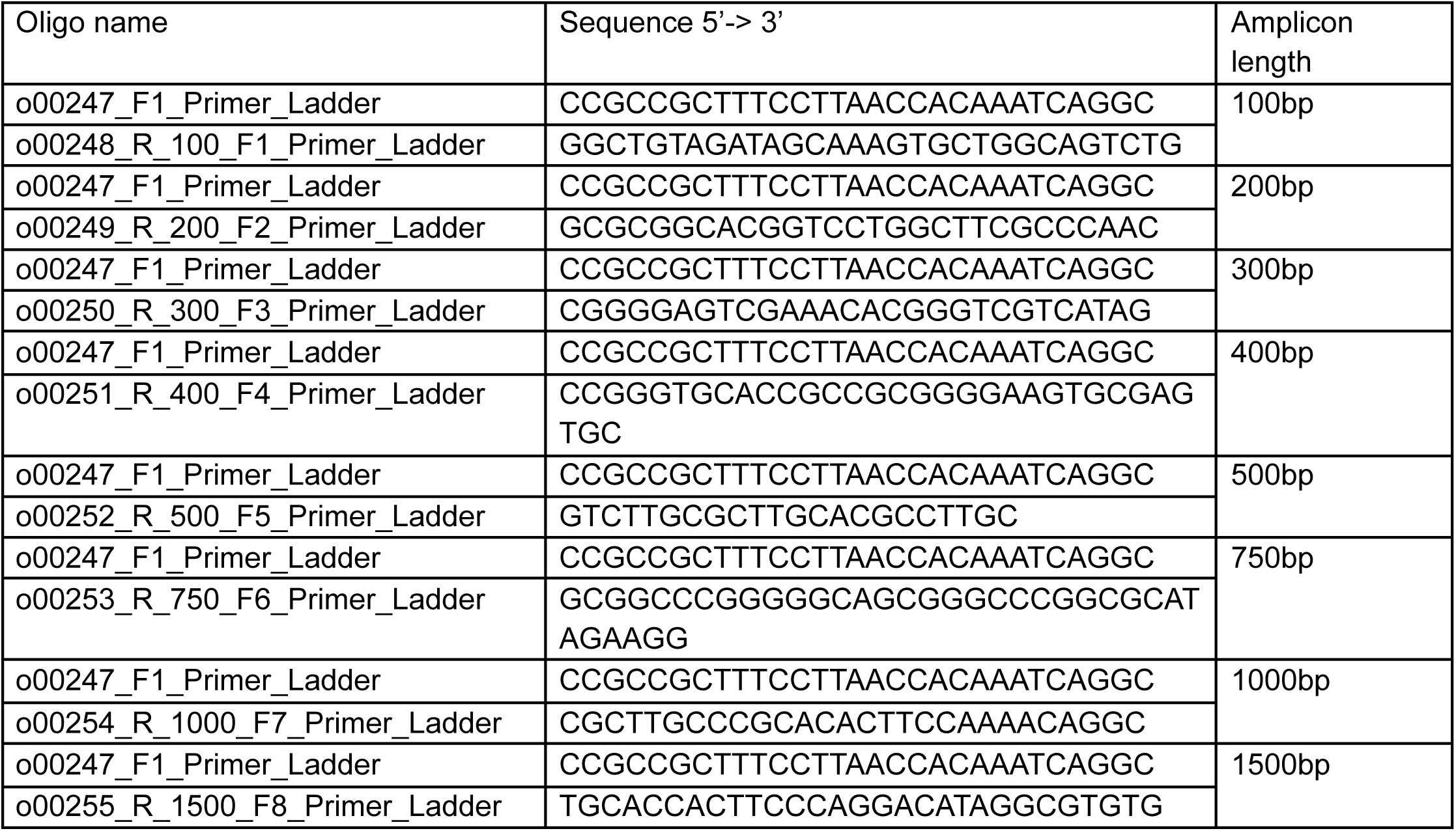

Supernatant was recovered by pelleting SPCs and stored. SPCs were washed 5x in 200µL Wash Buffer 2 (WB2) [10mM Tris/HCl pH 7.5, 0.1% Triton-X]. The same 0.5µM of the forward and reverse primer were then added to the corresponding SPCs in addition to Kapa HiFi 2x Master Mix (Roche; KK2602) in a 50µLreaction. After 30 cycles of a second PCR (same parameters), SPCs were washed 5x in 200µL Wash Buffer 1 (WB1) [10mM Tris/HCl pH 7.5, 0.1% Triton-X, 10mM EDTA]. SPCs were stained by adding 1µL of SYBR Green (200x working concentration) to pelleted SPCs with 100µL of 1x PBS. SPCs were imaged with a 4x objective by loading concentrated SPCs onto a countess slide. Gel electrophoresis was performed after breaking 5µL of SPCs by adding 1µL of Release Agent (Atrandi; CRP-PA1). Supernatant and SPC fractions were run on a 6% TBE gel, stained with SYBR Gold, and imaged.

### Primary Template Amplification (Figure S11)

Genomic DNA SPCs were generated by lysing HEK-293T cells in acid guanidinium thiocyanate cell extraction buffer (GeneJet RNA Purification Kit, Thermo Fisher) and washing twice in PBS. Unless indicated otherwise, 3µL of genomic DNA SPCs were provided as input for primary template amplification (ResolveDNA v2.0, BioSkryb Genomics). First, 1.68µL × L1, 0.12µL × L2, and 1.2µL × L3 reagents (buffer L123) were combined and 3µL of buffer L123 was added to SPCs and mixed at 1,400 RPM at RT for 20 minutes. Next, 5.4µL × R1, 0.6µL × R2 reagents (buffer R12) and optionally 0.56µL of SYBR-Green were combined and 6µL of buffer R12 was added to the reaction mixture and was mixed at 1,000 RPM at RT for 1 minute. A BioRad CFX Thermocycler was used with the following program optionally measuring SYBR-Green fluorescence every 30 s: 30°C for 105 min, 65°C for 3 min then stored at 4°C indefinitely.

Supernatant from the reaction mixtures was removed carefully before washing the SPCs 4 times with wash buffer containing 10mM Tris-HCl pH 8 (Invitrogen), 0.1% Triton-X (Sigma Aldrich), and 10mM EDTA pH 8.0 (Invitrogen). Where indicated, supernatant and extracted SPCs were run on a 1% (w/v%) agarose in TBE buffer.

### Amplicon data (Figure 3C-F)

SPCs containing K562 (ATCC; CCL-243) and PBMC (ATCC; PCS-200-011) cells were generated separately before mixing at different ratios. PCR targets were selected based on a previous report of K562 mutations (REF), and primer sequences along with expected amplicon lengths are provided in **Table S3** (Integrated DNA Technologies; standard desalting). Cell lysis and DNA denaturation were performed by treating SPCs with lysis buffer consisting of 400mM KOH, 10mM EDTA, 100mM DTT for 15 minutes, followed by five washes in 1M Tris-HCl with 0.1% Pluronic F-68 and five washes in Wash Buffer (10mM Tris-HCl pH 8.0, 0.1% v/v Pluronic F-68).

200µL PCR reactions combined 60µL of packed SPCs, 1x Platinum SuperFi II PCR Master Mix (Thermo Fisher Scientific; 12368010), and 0.5µM of each primer. Targets were amplified with the following program:

------------------------------------------------

98°C for 30 seconds

16 cycles of:

> 98°C for 20 seconds
>
> 60°C for 20 seconds
>
> 72°C for 1 minute

72°C for 5 minutes

------------------------------------------------

Two variations of further SPC-contained amplicon library prep were performed to obtain intact or fragmented amplicon data. Intact amplicon barcoding results in amplicon end sequencing data, whereas fragmentation prior to barcoding provides full amplicon length coverage. Both types of libraries were sequenced using a MiSeq Nano 300-cycle kit (Illumina; 15036522).

Intact amplicon barcoding and library prep: Following three SPC washes in Wash Buffer, amplicons were dA-tailed by combining 50µL of packed SPCs, 10µL of NEBuffer 2, 2µL of 10mM dATP, 6µL of Klenow fragment (NEB; M0212L), and 33µL of water, and incubating for 30 minutes at 37°C. Split and pool barcoding was performed using the commercially available Custom DNA Barcoding Module (Atrandi Biosciences; CKT-BARK4Q, User Guide DGPM02324192002). Four rounds of ligation-based combinatorial barcoding generated 24^4^ barcode combinations, and the first barcode was used to encode sample information (plate Col1, Col2, and Col3 for 100% K562, 20:80 K562:PBMC, and 10:90 K562:PBMC, respectively). After barcoding, DNA was released from SPCs using the Release Reagent (Atrandi Biosciences; CRP-RR1), and purified with 0.8x SPRI beads. Amplicons bearing a fully assembled barcode were enriched using a primer targeting a region upstream of the fully assembled barcode (p7_PCR_ix2, **Table S3**) and the reverse primers used during the initial multiplex PCR. Fragmentation and second Illumina adapter ligation were performed as described in User Guide DGPM02323206001 (Atrandi Biosciences). A 100µL final indexing PCR was composed of 40µL purified DNA, 5 µM p7_PCR1_ix2, 5µM p5_PCR (**Table S3**), and 1x Platinum SuperFI II PCR Master Mix. PCR was performed with the program below and purified with 0.6x-0.8x SPRI beads:

------------------------------------------------

98°C for 30 seconds

16 cycles of:

> 98°C for 20 seconds
>
> 60°C for 30 seconds
>
> 72°C for 1 minute

72°C for 5 minutes

------------------------------------------------

Fragmented amplicon barcoding library prep: Following three SPC washes in Wash Buffer, amplicon fragmentation, end-prep and A-tailing were performed by combining on ice 50µl of packed SPCs, 20µl of NEBNext Ultra II FS Reaction Buffer, 5.72µl NEBNext Ultra II FS Enzyme Mix, 24.3µl of water, and incubating for 10 min at 37°C followed by 30 min at 65°C. Following 5 washes in Wash Buffer, further split and pool barcoding was performed using the Custom DNA Barcoding Module (Atrandi Biosciences; CKT-BARK4Q, User Guide DGPM02324192002) with a modification. Specifically, 2µl of 100 µM Ligation Adapter (composed of pre-annealed /5Phos/GATCGGAAGAGCGTCGTGTAGGGAAAGAGTG*T and /5AmMC6/GCTCTTCCGATCT; HPLC purification) were spiked in each of the 24 barcode A wells, i.e. columns 1-3 of the Barcode plate (Atrandi Biosciences; CRP-PLT1). After barcoding, DNA was released from SPCs using the Release Reagent, and purified with 0.8x SPRI beads, directly followed by a 100µL indexing PCR composed of 28µL purified DNA, 5 µM p7_PCR1_ix1, 5µM p5_PCR (**Table S3**), and 1x Platinum SuperFI II PCR Master Mix. PCR followed the same program as for intact amplicons but for 20 cycles, followed by 0.6-0.8x SPRI clean-up.

**Supplemental Table 3:**
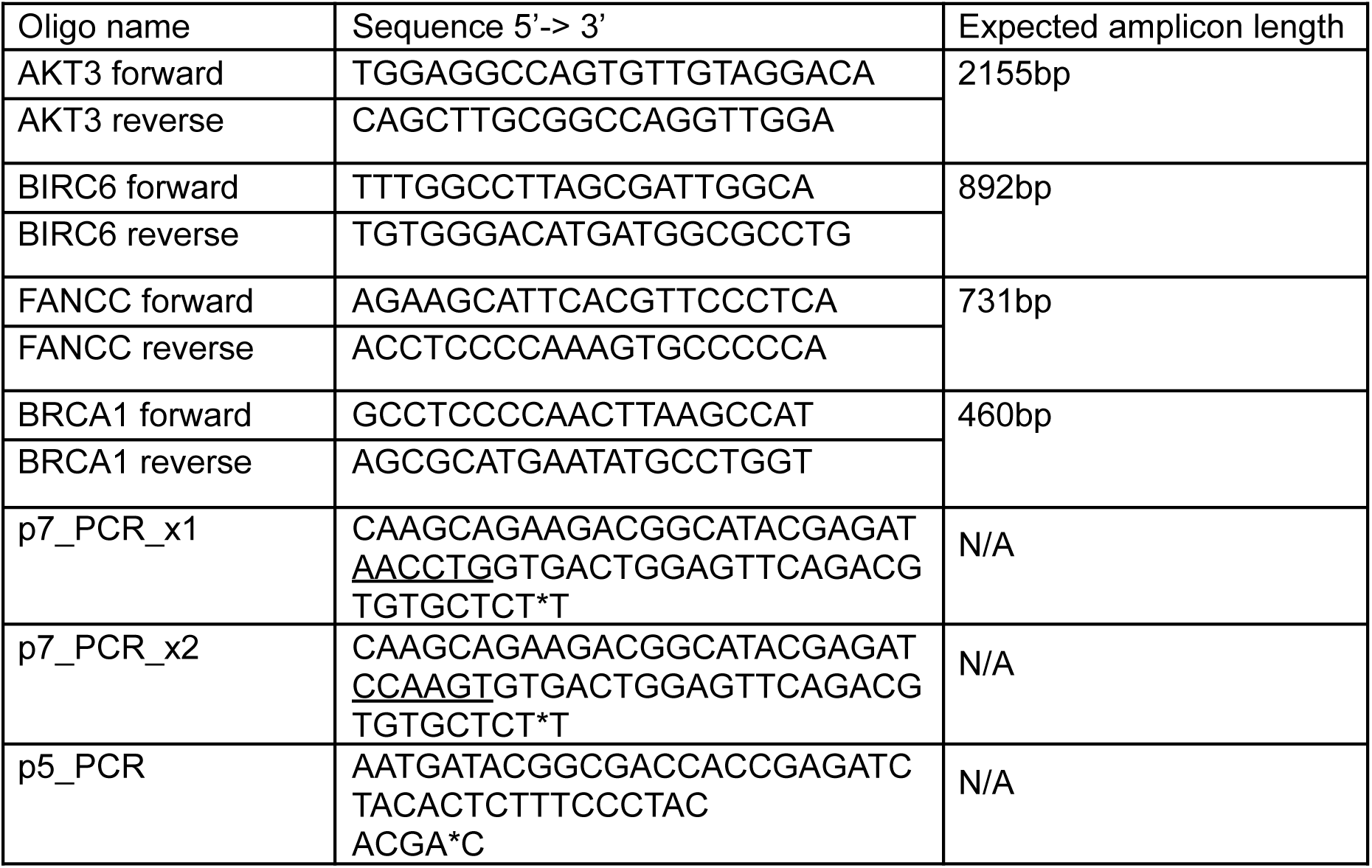

### Amplicon data analysis (Figure 3C-F)

Raw sequencing reads were demultiplexed and aligned to hg38 as described in preprocessing (below). Cells with low coverage (read counts lower than 100) were removed. Variant calling was performed withGATK4 (*46*). Reads are grouped by cell labels and deduplication is run on each individual cell. Haplotype caller was used to generate GVCFs for each individual cell before joint calling. The default quality score threshold of 10 was used for filtering mutations.

### Sequencing Data Preprocessing (Figure 3G-I, Fig. S13)

Raw illumina bcl files were converted to fastq format using illumina bcl2fastq (version 2.20.0). A custom python script was used to extract the individual cell barcodes from read 2 and perform error correction of the barcodes, based upon a calculated maximum permissible Levenshtein distance given the combination of oligos used. Reads whose barcodes exceeded the maximum distance were discarded prior to alignment. Sequencing adapters were trimmed from reads using Trim Galore (version 0.6.10). Reads were then aligned to the reference genome using bwa-mem (version 0.7.17) (*47*). For the species mixing experiments, a hybrid human-mouse reference genome was created by concatenating HG38 and MM10, and the number of reads aligning to either the mouse or human genomes was computed for each cell. Barcodes associated with empty SPCs were differentiated from barcodes associated with occupied SPCs by ranking barcodes by number of associated reads and plotting aligned read counts per barcode. Next, a cutoff was either selected manually based upon the anticipated number of captured cells and the shape of the plot, or by algorithmically estimating the number of recovered cells based upon the known capture rate with sub-Poisson loading. Variant calling was performed with either bcftools call (for low coverage genome data) (*48*) or GATK haplotype caller (for higher coverage amplicon data) (*46*). For bulk WGS data, the -m flag was used for multiallelic calling and resulting bcf files were standardized using the bcftools norm command.

### Barnyard Whole Genome Amplification Experiment (Figure 3G-I, Fig. S13)

SPCs were generated with the standard protocol by encapsulating a mixture of 125,000 NIH3T3 cells and 125,000 HEK293T cells. After capsule polymerization, emulsion breaking and a wash in 1x PBS, capsules were suspended in 1000µL of ice-cold methanol and stored at -20C. 250µL of hard packed SPCs (500µL of a 1:1 mixture of methanol and SPCs) were then transferred to a new 1.5-mL low-bind Eppendorf tube. These SPCs were first pelleted (2000g for 30 seconds in a cooled bench top centrifuge) and then the methanol was removed and replaced with 1 mL of Wash Buffer 2 (WB2: 10mM Tris/HCl pH 7.5, 0.1% Triton-X). This was followed by 4 more washes with WB2. After the 5th wash, SPCs were resuspended to a final volume of 1 mL in WB2 10µL of RNaseA (Thermo Scientific Scientific; R1253) was added to the SPCs and the SPCs were rotated end over end for 45 minutes at 37°C. After incubation SPCs were spun down (2000g for 30 seconds) and the buffer was then replaced with Wash Buffer 1 (WB1: 10mM Tris/HCl pH 7.5, 0.1% Triton-X, 10mM EDTA). After 4 more washes in WB1, SPCs were resuspended in WB2 and washed 10 times. At this point modified MDA or PTA was performed on SPCs.

Modified MDA: Using the Single-Microbe DNA Barcoding Kit (Atrandi Biosciences; CKP-BARK1), SPCs were washed in 1 mL of 1x WB (Atrandi Biosciences; CRP-WB1). Then SPCs were resuspended in 1 mL of 1:1 dilution of Lysis Buffer (Atrandi Biosciences; CRP-LB1) supplemented with 100µL of 1M DTT. These SPCs were incubated for 15 minutes at room temperature with end over end rotation to denature DNA. These SPCs were then pelleted, the supernatant was aspirated and the reaction was neutralized with five 1-mL washes with buffer Neutralization Buffer (Atrandi Biosciences; CRP-NB1). This was followed by five additional 1-mL washes in buffer WB. After the final wash the volume of SPCs was adjusted to 150µL with distilled water and the following components were added: 97.5µL nuclease free water, 37.5µL 10x WGA Reaction Buffer (Atrandi Biosciences; CRP-WGB1), 37.5µL dNTP mix (Atrandi Biosciences; CRP-DNTP1), 18.75µL of Primer Mix (Atrandi Biosciences; CRP-PRM1), 3.75µL of 0.1M DTT, 3.75µL 10% Pluronic F-68, and 7.5µL of WGA Enhancer (Atrandi Biosciences; CRP-WGE1). This mixture was then heated to 95°C for 5 minutes on a thermomixer to allow primers to anneal and snap cooled on ice. After cooling 18.75µL of WGA polymerase (Atrandi Biosciences; CRP-WGP1) was added to the reaction mixture and allowed to incubate for 15 minutes at 45°C on a thermomixer followed by an incubation at 65°C for 10 minutes. After amplification, SPCs were pelleted and washed 3x with 1 mL of WB.

PTA: PTA was performed using the Bioskryb Resolve v2 kit (Bioskryb Genomics; PN 100545) following the manual TAS-073. SPCs were first washed 5x with 1x PBS. Followed by a wash with 1 volume of Cell Wash Buffer (Bioskryb Genomics; PN 100002) – 40µL of closely packed SPCs was removed and added to 40µL of Cell wash buffer. This wash was performed a total of 3 times. After the final wash, SPCs were pelleted at 2000g for 1 minute and excess supernatant was aspirated. Initial reaction was performed using 40µL of input SPC template with 6.72µL of L1 reagent (Bioskryb Genomics; PN 100522), 0.48µL L2 Reagent (Bioskryb Genomics; PN 100016), and 4.8µL L3 reagent (Bioskryb Genomics; PN 100523). This mixture was allowed to mix end-over-end for 20 minutes at room temperature. After incubation, 21.6µL of R1 (Bioskryb Genomics; PN 100521) and 2.4µL of R2 (Bioskryb Genomics; PN 100527) were added and mixed before incubation in a thermocycler set to 30°C for 2.5 hours followed by a 5 minute incubation at 65°C. After amplification SPCs were pelleted and washed 3x with 1 mL of WB.

Combinatorial indexing: After washing WGA SPCs in WB, a small aliquot of SPCs was imaged to ensure DNA amplification. After a 30-minute end-prep reaction (Atrandi Biosciences; CRP-EPE1, CRP-EPB1), SPCs from the two WGA conditions were loaded into a distinct set of wells. For mMDA, Col1 (A-H) and Col2 (A-G) were used from the barcoding module (Atrandi Biosciences; CRP-PLT1) and Col2 (H) and Col3 (A-H) were used for PTA reactions (Atrandi Biosciences; CRP-PLT1). The SPCs were resuspended with 60μL of 10x Ligation Buffer (Atrandi Biosciences; CRP-LGB1) and 20µL of Ligation Enzyme (Atrandi Biosciences; CRP-LGE1) and brought to a final volume of 300µL with distilled water. Ligation was performed for 15 minutes at room temperature on a plate shaker. After ligation, 40µL of Stop Buffer (Atrandi Biosciences; CRP-SB1) was added to each well and the mixture was allowed to incubate for 5 minutes at room temperature. After ligation, SPCs were washed 5x with WB and resuspended in ligation mix again before redistribution to the next set of barcoding wells. This process was repeated for a total of 4 rounds. After the final ligation SPCs were stained with SYBR green (Thermo Fisher Scientific; S7563) and fluorescent SPCs were counted on a hemocytometer. Fluorescent DNA signal was present in 70 SPCs/μL with a total SPC solution volume of 1500µL. This indicated that 105,000 SPCs containing amplified DNA were present at this point.

Index PCR: 10µL of SPCs with an estimated 700 cells or 25µL of SPCs with an estimated 1,750 cells were removed from the pool and distilled water was added to each to bring the volume up to 50µL. To these solutions, 1µL of Release Reagent (Atrandi Biosciences; CRP-RR1) was added and the SPCs were allowed to incubate at room temperature for 5 minutes until fully released. Next 40µL of room temperature SPRI beads (Beckman Coulter; B23319) was added to the released SPC solution. The bound fraction was eluted in 27µL of distilled water and 1µL of the eluate was quantified using the Qubit High Sensitivity Assay kit (Thermo Fisher Scientific; Q32851). 7µL of 5x Fragmentase buffer (NEB; B0349) and 2µL of Fragmentase (NEB; M0348) were added to 26µL of eluted DNA on ice. Reaction was allowed to proceed for 10 minutes on a warmed thermocycler at 37°C before incubation at 65°C for 30 minutes. To this sample, 30µL of Ligation Master Mix (NEB; E7648) was added in addition to 1µL of Ligation Enhancer (NEB; E7374) and 2.5µL of Ligation Adapter (Atrandi Biosciences; CRP-LGA1). This reaction was mixed and allowed to incubate at 20°C for 15 minutes before a 0.8x SPRI cleanup was performed. The DNA was eluted in 40µL and added to a mastermix containing 50µL of Q5 Ultra Master Mix (NEB; M0544), 10µL of primers (Atrandi Biosciences; CRT-PRM2) and 0.5µL of 200x SYBR Green. PCR was performed with the following program:

------------------------------------------------

98°C for 45 seconds

15 cycles of:

> 98°C for 20 seconds
>
> 54°C for 30 seconds
>
> 72°C for 20 seconds

72°C for 2 minutes

------------------------------------------------

Libraries were then cleaned up using a 2-sided SPRI cleanup (0.5x, 0.7x), quantified via Qubit High Sensitivity Assay and visualized on the Tapestation D1000HS (Agilent Technologies).

Sequencing: These libraries were loaded on a P2 200 cycle NextSeq2000 kit (Illumina) at a loading concentration of 800 pM with the following read structure: Read1:82, Index1:10, Index2:10, Read2:107 without custom sequencing primers.

### Estimation of coverage uniformity and library complexity (Figure 3G-I, Fig. S13)

GATK EstimateLibraryComplexity was used to estimate the number of unique molecules in the sequencing libraries. To compare coverage uniformity between amplification techniques, coverage track bigwig files with 1-kilobase bins were generated using deepTools bamCoverage. A custom Python script was used to generate a Lorenz curve, depicting the distribution of read depth across regions of the genome for each cell. To account for regions lacking coverage, bins with zero coverage were excluded. Bins were sorted in ascending order by read depth, and the cumulative proportion of total reads across the bins was plotted against the cumulative proportion of bins. The Gini coefficient (*G*), defined as *G* = 1 − 2*A* where *A* is the area under the Lorenz curve, was computed for each cell by approximating the area under the lorenz curve using the trapezoidal rule with NumPy’s trapz function.

### POLE P286R editing (Figure 4)

To generate K562 POLE^P286R^ cells, we used CRISPR-Cas9 to generate a double-stranded DNA break around that locus and introduced a homology-directed repair (HDR) template carrying that specific amino acid substitution. For each experimental group, up to 5 x 10^6^ cells were washed three times in cold DPBS (Gibco) and resuspended in supplemented SF solution (Lonza) at a concentration of 1 x 10^6^ cells per 10µL. The cells were combined with 10μg of Cas9 (QB3 MacroLab), 5μg of sgRNA (IDT), 3.3μg of HDR template (IDT), and 16.8pmol of electroporation enhancer (IDT). Nucleofection was performed using the 4D-Nucleofector X unit (Lonza) using the specified program for K562 cells. After nucleofection, cells were returned to pre-warmed media supplemented with 1µM Alt-R HDR Enhancer V2 (IDT).

### POLE P286R genotyping PCR (Figure 4, Figure S14)

To quantify initial editing rates, we isolated gDNA, performed targeted PCR at the editing locus, sequenced the amplicons, and measured the percentage of reads with the specific nucleotide substitution encoding POLE P286R using CRISPResso2. gDNA was isolated either using the DNeasy Blood & Tissue kit (Qiagen) per the manufacturer’s protocol or lab-prepared lysis buffer containing proteinase-K [lysis buffer: 10mM Tris/HCl, 0.05% SDS, 0.1mg/mL Proteinase-K (Thermo Fisher; EO0492)]. Pelleted cells were resuspended in lysis buffer for 2 hours at 37°Cbefore heat inactivation for 30 minutes at 80C. For gDNA isolated with the DNeasy Blood & Tissue kit, targeted amplification of the POLE editing locus was performed using the primers o00230_P286R_PCR_F and o00231_P286R_PCR_R. For gDNA isolated with lab-prepared lysis buffer, targeted amplification was performed using the primers o00312_PolE_P286R_genome_F and o00313_PolE_P286R_genome_R. Amplicons were PCR cleaned using either the QIAquick PCR Purification kit (Qiagen) or a 0.8x right sided SPRIselect (Beckman Coulter) bead cleanup. Purified amplicons were sent to Plasmidsaurus for premium PCR sequencing and quantified using CRISPResso2 in batch mode using the following parameters:

--batch_settings batch.bat

--amplicon_seq ATGGGGAGTTTAGAGCTTGGCTTTATGCTTATTTTGTCCCCACAGGACCCTGTGGTTTTGGCATTTGACATTGAGACGACCAAACTGCCCCTCAAGTTTCCTGATGCTGAGACAGACCAGATTATGATGATTTCCTACATGATCGATGGCCAGGTG

-p 8

-n nhej

-g ATCTGGTCTGTCTCAGCATC

-wc 0

-w 5

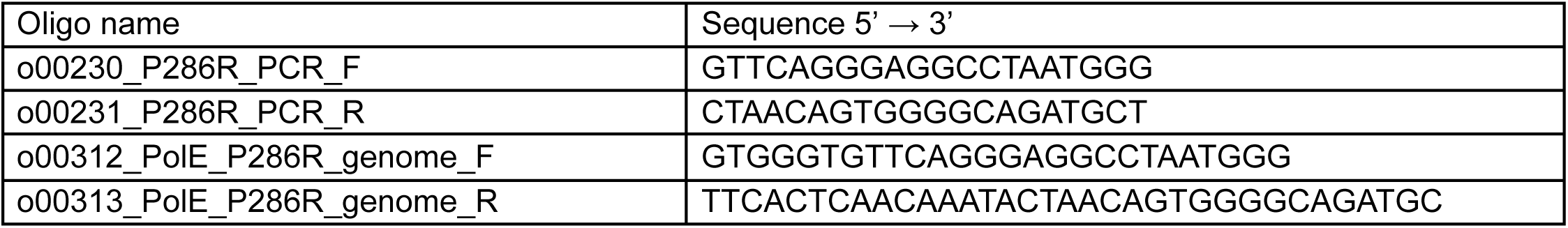

### POLE P286R bulk whole genome sequencing (Figure 4B)

Healthy and cycling K562 cells were nucleofected with CRISPR-Cas9 RNP on Day 0. Cells were counted on days 2, 4, 6, 8, 10, 12, and 14 and provided growth media to maintain a concentration of approximately 1,000 cells perµL. On days 3, 6, 9, 12 and 15, genomic DNA from between 1.5 x 10^5^ and 1 x 10^6^ of cycling cells was collected by pelleting cells. Cell pellets were frozen and stored at -20°Cuntil they were processed further.

All samples were thawed, resuspended in lysis buffer containing proteinase-K [Lysis buffer: 10mM Tris/HCl, 0.05% SDS, 0.1mg/mL Proteinase-K (Thermo Fisher; EO0492)] for 2 hours at 37C. The buffer was then exchanged via a 1x SPRI cleanup (Beckman Coulter; B23319) and the eluted DNA was quantified using the Qubit High Sensitivity kit. 100ng of each sample was then prepared for sequencing using the NEBNext® Ultra II FS library preparation kit (NEB; E7805) with a slight modification – each of the reaction volumes was reduced to a third of the recommended volume. During index PCR libraries were amplified for 10 cycles before pooling and sequencing. Libraries were sequenced on a NovaSeqX 300 cycle kit at a loading concentration of 750pM.

### Hypermutator Cell Line Characterization (Figure 4B)

A modified version (https://github.com/dustin-mullaney/mutyper_rare_variant) of the Mutyper python package (*49*) was used to quantify the number of SNPs and characterize their trinucleotide context in bulk and single cell genome datasets. In bulk samples, only sites with at least ten reads were considered, and they were only called as a SNP if at least two reads supported the alt allele. To compute the variant allele frequency and make comparisons between samples with uneven sequencing depth, scipy.stats (version 1.15.2) was used to downsample sites exceeding the depth threshold of ten reads with the hypergeometric distribution. Counts of mutations were normalized between experimental conditions by the total number of reads for each condition. Statistical analysis and linear regression was conducted with scipy.statsmodels (version 0.14.4).

### POLE P286R Clonal Bottlenecking (Figure 4C-G)

Previous attempts to make single cell colonies of K562 cells resulted in cellular death. To get viable bottlenecked colonies approximately 1,000 edited K562 cells were plated via dilution into 2 round bottom (Thermo Scientific; 163320) 96 well plates with 100μL of growth media 1 day after transfection with the CRISPR Cas9 gRNA RNP. These colonies were monitored daily, and on Day 8 post transfection, six colonies were moved to a new round bottom plate and split into two wells. Genomic DNA from one of the wells was used to assess the fidelity of editing at the POLE P286R allele. The other well was split in the following fashion (diagrammed in **Figure 4C**): On Day 9 the well was split 1:1 to a new well; on day 11 one of the day 9 wells was split 1:1 and so on. This pattern was repeated on days 13, 15, 17, and 19. Out of the six clones selected 3 displayed editing at the locus. The wells from the chosen clone (Clone F5) were counted and encapsulated for scWGS in SPCs. The other clones were frozen in freezing media [90% FBS (v/v, 10% DMSO (v/v)] and stored in liquid nitrogen.

### Whole Genome Amplification and combinatorial indexing of clonally restricted K562s (Figure 4C-G, Fig. S15)

Encapsulation: Six individual splits (see **Figure 4C** for reference) were encapsulated separately. Standard encapsulation procedure was followed using the 65µm SPC kit (Atrandi Biosciences; CKN-G11). Cells were counted and diluted to 1x 10^7^ cells/mL and 1.5 x 10^5^ (15µL) of cells was added to the working core solution. SPCs were collected, and polymerized. Emulsions were broken and capsules were dehydrated through the addition of 1 mL of 100% methanol and stored at -20°C until further processing.

SPC cell lysis: All spins were at 2000g for 30 seconds at 4°C. First, SPCs were vortexed to break clumps and resuspend. 300µL of SPCs stored in methanol were then removed and moved to a new tube. SPCs were spun down and methanol was removed by aspiration. SPCs were resuspended in 800µL of freshly prepared lysis buffer [10 mM Tris/HCl, 0.05% SDS]. Lysis buffer was added in a dropwise manner with slight agitation to rehydrate. SPCs were then allowed to mix end over end for 2 minutes before an additional spin down and wash with another 800µL of lysis buffer. SPCs were then spun down and the supernatant was aspirated. Additional 500µL of lysis buffer was added to each batch of SPCs along with 20µL of Proteinase K (Thermo Fisher Scientific; EO0492). This mixture was set on the thermomixer for 1 hour and proteins were allowed to digest at 37ľC with the shaker set to 1000 rpm. A 1-μL aliquot of each was stained with SYBR Green and imaged to check that the nucleus was sufficiently digested. After digestion, SPCs were pelleted, the supernatant was removed and the SPCs were washed washed 2x with 800µL of Wash Buffer 1 (WB1) [10mM Tris/HCl pH 7.5, 0.1% Triton-X, 10mM EDTA].

Whole Genome Amplification: Resolve Whole Genome Amplification materials (Bioskryb Genomics; PN 100545) PTA reagents were allowed to thaw at room temperature or on ice. Each batch of SPCs was washed 2x using 1 mL of 1x PBS to remove EDTA residual from WB1. 50µL of closely packed SPCs were removed from each sample and 100µL of Cell wash buffer (Bioskryb Genomics; PN 100002) was used to wash each sample. This wash was repeated 2x for a total of 3 washes. After the final wash, the following mix was made: 107.25µL L1 (Bioskryb Genomics; PN 100522), 7.8µL L2 (Bioskryb Genomics; PN 100016) and 78µL L3 (Bioskryb Genomics; PN 100523). 30µL of this master mix was added to approximately 50µL of SPCs in each condition. This mixture was allowed to mix end-over-end for 20 minutes at room temperature. A scaled reaction master mix was made containing 351µL R1 (Bioskryb Genomics; PN 100521) and 39µL R2 (Bioskryb Genomics; PN 100527). After 20 minutes, 60µL of the reaction master mix was added to each tube of SPCs. SPCs were mixed on a thermomixer set at 37°C and 1000 rpm for 2.5 hours and heat inactivated at 65°C for 5 minutes after amplification. All 6 samples were washed 5x in Wash Buffer 2 (WB2) [10mM Tris/HCl pH 7.5, 0.1% Triton-X].

Fragmentation, and combinatorial indexing: Next using the FS II Ultra Fragmentation module, 30µL of SPCs were mixed with 8µL of NEB Next Ultra II Reaction Buffer (NEB; B0349) and 2.26µL of NEB Next Ultra II FS Enzyme (NEB; M0348) and allowed to incubate with the following program: 37°C for 7.5 minutes (fragmentation), followed by 65°C for 30 minutes (A-tailing and end repair). SPCs were then washed 3x with Wash Buffer WB (Atrandi Biosciences; CRP-WB1) and combinatorial indexing was performed. Samples were placed in specific wells of the first combinatorial indexing reaction as follows from the barcoding module (Atrandi Biosciences; CKT-BARK4Q) of the Single-Microbe DNA Barcoding kit (CKP-BARK1): P5:Col1 A-D; P4:Col1 E-H; P3:Col2 A-D; P2:Col2 E-H; P1:Col3 A-D and P0:Col3 E-H. The SPCs were resuspended with 10µL of 10x Ligation Buffer (Atrandi Biosciences; CRP-LGB1) and 3.33µL of Ligation Enzyme (Atrandi Biosciences; CRP-LGE1) and brought to a final volume of 50µL with distilled water. 10µL of SPC:ligation mix was added to each well of the barcoding reaction. Ligation was performed for 15 minutes at room temperature on a plate shaker. After ligation, Stop Buffer (Atrandi Biosciences; CRP-SB1) was added to each well and the reaction was allowed to incubate for 5 minutes at room temperature. After ligation, SPCs were washed 5x with Wash Buffer and resuspended in the ligation mix again before redistribution to the next set of barcoding wells. This process was repeated for a total of 4 rounds. After the final ligation SPCs were stained with SYBR green and fluorescent SPCs were counted on a hemocytometer. Fluorescent DNA signal was present in 33 DNA filled SPCs/μL of the sample.

Index PCR: 33µL of SPCs corresponding to an estimated 1000 cell-containing SPCs were removed from the pool and distilled water was added to bring the volume up to 50µL. To this solution, 1µL of release agent (Atrandi Biosciences; CRP-RR1) was added and the SPCs were allowed to incubate at room temperature for 5 minutes until fully released. Next, 40µL of room temperature SPRI beads (Beckman Coulter; B23319) was added to the released SPC solution. The bound fraction was eluted in 27µL of distilled water and 1µL of the reaction was quantified using the Qubit High Sensitivity Assay kit (Thermo Fisher Scientific; Q32851). 7µL of 5x Fragmentase buffer (NEB; B0349) and 2µL of Fragmentase (NEB; M0348) were added to 26µL of eluted DNA on ice. Reaction was allowed to proceed for 9 minutes on a warmed thermocycler at 37°C before incubation at 65°Cfor 30 minutes. To this sample, 30μL of Ligation Master Mix (NEB; E7648) was added in addition to 1µL of Ligation Enhancer (NEB; E7374) and 2.5µL of Ligation Adapter (Atrandi Biosciences; CRP-LGA1). This reaction was mixed and allowed to incubate at 20°C for 15 minutes before a 0.8x SPRI cleanup was performed. A second 0.8x SPRI cleanup was performed and DNA was eluted into 160µL of distilled water. This purified DNA was added to a mastermix containing 200µL Q5 Ultra Master Mix (NEB; M0544), 40µL of primers (Atrandi; CRT-PRM2) and 2µL of 200x SYBR Green. PCR was performed with the following program:

------------------------------------------------

98°C for 45 seconds

15 cycles of:

> 98°C for 20 seconds
>
> 54°C for 30 seconds
>
> 72°C for 20 seconds

72°C for 2 minutes

------------------------------------------------

Libraries were then cleaned up using a 2-sided SPRI cleanup (0.5x, 0.75x), quantified via Qubit High Sensitivity Assay and visualized on the Tapestation D1000HS (Agilent Technologies).

Sequencing: These libraries were loaded on a P3 300 cycle NextSeq2000 kit (Illumina) at a loading concentration of 750 pM with the following read structure: Read1:150, Index1:10, Index2:10, Read2:150 without custom sequencing primers.

### Lineage Tracing (Figure 4C-G)

Following demultiplexing and alignment, variants were called on both the bulked dataset and across all individual cells, compared to hg38. For downstream analysis, the resulting data was consolidated into an anndata object (*50*) with three layers, each indexed by cells (.obs_names) and variant sites (.var_names): the primary layer contains a boolean matrix which indicates the presence or absence of *any* reads supporting the alternative allele at a given site, a second layer stores the read depth for each cell at each site, and the third layer stores the total number of reads supporting the alternative allele for each cell at each site. Any sites that were called as homozygous or heterozygous alt in the bulked dataset were considered to be K562 germline variants, and were removed from the anndata object, such that only de novo SNPs caused by POLE P286R were considered for lineage tracing. For grouped lineage tree construction, variants were aggregated within populations, and hamming distances were computed between each population to create a distance matrix. The unweighted pair group method with arithmetic mean (UPGMA) was used to construct a phylogenetic tree of the aggregated cells. For single cell lineage tracing, hamming distances were computed between pairings of individual cells by considering all sites that were observed in both cells (more than two reads in each) and the tree was constructed with the UPGMA algorithm. For both single cell and grouped lineage tracing, the hamming distances are normalized by the number of sites considered.

## References

1. A. Tanay, A. Regev, Scaling single-cell genomics from phenomenology to mechanism. Nature 541, 331–338 (2017).

2. A. K. Shalek, R. Satija, X. Adiconis, R. S. Gertner, J. T. Gaublomme, R. Raychowdhury, S. Schwartz, N. Yosef, C. Malboeuf, D. Lu, J. J. Trombetta, D. Gennert, A. Gnirke, A. Goren, N. Hacohen, J. Z. Levin, H. Park, A. Regev, Single-cell transcriptomics reveals bimodality in expression and splicing in immune cells. Nature 498, 236–240 (2013).

3. P. Datlinger, A. F. Rendeiro, C. Schmidl, T. Krausgruber, P. Traxler, J. Klughammer, L. C. Schuster, A. Kuchler, D. Alpar, C. Bock, Pooled CRISPR screening with single-cell transcriptome readout. Nat Methods 14, 297–301 (2017).

4. A. Dixit, O. Parnas, B. Li, J. Chen, C. P. Fulco, L. Jerby-Arnon, N. D. Marjanovic, D. Dionne, T. Burks, R. Raychowdhury, B. Adamson, T. M. Norman, E. S. Lander, J. S. Weissman, N. Friedman, A. Regev, Perturb-Seq: Dissecting Molecular Circuits with Scalable Single-Cell RNA Profiling of Pooled Genetic Screens. Cell 167, 1853–1866.e17 (2016).

5. A. M. Klein, L. Mazutis, I. Akartuna, N. Tallapragada, A. Veres, V. Li, L. Peshkin, D. A. Weitz, M. W. Kirschner, Droplet barcoding for single-cell transcriptomics applied to embryonic stem cells. Cell 161, 1187–1201 (2015).

6. E. J. Walsh, A. Feuerborn, J. H. R. Wheeler, A. N. Tan, W. M. Durham, K. R. Foster, P. R. Cook, Microfluidics with fluid walls. Nat Commun 8, 816 (2017).

7. E. Brouzes, M. Medkova, N. Savenelli, D. Marran, M. Twardowski, J. B. Hutchison, J. M. Rothberg, D. R. Link, N. Perrimon, M. L. Samuels, Droplet microfluidic technology for single-cell high-throughput screening. Proc Natl Acad Sci U S A 106, 14195–14200 (2009).

8. S. Köster, F. E. Angilè, H. Duan, J. J. Agresti, A. Wintner, C. Schmitz, A. C. Rowat, C. A. Merten, D. Pisignano, A. D. Griffiths, D. A. Weitz, Drop-based microfluidic devices for encapsulation of single cells. Lab Chip 8, 1110–1115 (2008).

9. G. Leonaviciene, K. Leonavicius, R. Meskys, L. Mazutis, Multi-step processing of single cells using semi-permeable capsules. Lab Chip 20, 4052–4062 (2020).

10. G. Leonaviciene, L. Mazutis, RNA cytometry of single-cells using semi-permeable microcapsules. Nucleic Acids Res 51, e2 (2023).

11. J. L. Watson, D. Juergens, N. R. Bennett, B. L. Trippe, J. Yim, H. E. Eisenach, W. Ahern, A. J. Borst, R. J. Ragotte, L. F. Milles, B. I. M. Wicky, N. Hanikel, S. J. Pellock, A. Courbet, W. Sheffler, J. Wang, P. Venkatesh, I. Sappington, S. V. Torres, A. Lauko, V. De Bortoli, E. Mathieu, S. Ovchinnikov, R. Barzilay, T. S. Jaakkola, F. DiMaio, M. Baek, D. Baker, De novo design of protein structure and function with RFdiffusion. Nature 620, 1089–1100 (2023).

12. D. A. Cusanovich, R. Daza, A. Adey, H. A. Pliner, L. Christiansen, K. L. Gunderson, F. J. Steemers, C. Trapnell, J. Shendure, Multiplex single cell profiling of chromatin accessibility by combinatorial cellular indexing. Science 348, 910–914 (2015).

13. R. M. Mulqueen, D. Pokholok, B. L. O’Connell, C. A. Thornton, F. Zhang, B. J. O’Roak, J. Link, G. G. Yardımcı, R. C. Sears, F. J. Steemers, A. C. Adey, High-content single-cell combinatorial indexing. Nat Biotechnol 39, 1574–1580 (2021).

14. J. Cao, J. S. Packer, V. Ramani, D. A. Cusanovich, C. Huynh, R. Daza, X. Qiu, C. Lee, S. N. Furlan, F. J. Steemers, A. Adey, R. H. Waterston, C. Trapnell, J. Shendure, Comprehensive single-cell transcriptional profiling of a multicellular organism. Science 357, 661–667 (2017).

15. S. Li, J. Kendall, S. Park, Z. Wang, J. Alexander, A. Moffitt, N. Ranade, C. Danyko, B. Gegenhuber, S. Fischer, B. D. Robinson, H. Lepor, J. Tollkuhn, J. Gillis, E. Brouzes, A. Krasnitz, D. Levy, M. Wigler, Copolymerization of single-cell nucleic acids into balls of acrylamide gel. Genome Res. 30, 49–61 (2020).

16. A. B. Rosenberg, C. M. Roco, R. A. Muscat, A. Kuchina, P. Sample, Z. Yao, L. T. Graybuck, D. J. Peeler, S. Mukherjee, W. Chen, S. H. Pun, D. L. Sellers, B. Tasic, G. Seelig, Single-cell profiling of the developing mouse brain and spinal cord with split-pool barcoding. Science 360, 176–182 (2018).

17. A. Tsherniak, F. Vazquez, P. G. Montgomery, B. A. Weir, G. Kryukov, G. S. Cowley, S. Gill, W. F. Harrington, S. Pantel, J. M. Krill-Burger, R. M. Meyers, L. Ali, A. Goodale, Y. Lee, G. Jiang, J. Hsiao, W. F. J. Gerath, S. Howell, E. Merkel, M. Ghandi, L. A. Garraway, D. E. Root, T. R. Golub, J. S. Boehm, W. C. Hahn, Defining a Cancer Dependency Map. Cell 170, 564–576.e16 (2017).

18. Y. Wang, J. Waters, M. L. Leung, A. Unruh, W. Roh, X. Shi, K. Chen, P. Scheet, S. Vattathil, H. Liang, A. Multani, H. Zhang, R. Zhao, F. Michor, F. Meric-Bernstam, N. E. Navin, Clonal evolution in breast cancer revealed by single nucleus genome sequencing. Nature 512, 155–160 (2014).

19. T. J. Pugh, A. D. Delaney, N. Farnoud, S. Flibotte, M. Griffith, H. I. Li, H. Qian, P. Farinha, R. D. Gascoyne, M. A. Marra, Impact of whole genome amplification on analysis of copy number variants. Nucleic Acids Res 36, e80 (2008).

20. V. Gonzalez-Pena, S. Natarajan, Y. Xia, D. Klein, R. Carter, Y. Pang, B. Shaner, K. Annu, D. Putnam, W. Chen, J. Connelly, S. Pruett-Miller, X. Chen, J. Easton, C. Gawad, Accurate genomic variant detection in single cells with primary template-directed amplification. Proc Natl Acad Sci U S A 118 (2021).

21. J. Ganz, L. J. Luquette, S. Bizzotto, M. B. Miller, Z. Zhou, C. L. Bohrson, H. Jin, A. V. Tran, V. V. Viswanadham, G. McDonough, K. Brown, Y. Chahine, B. Chhouk, A. Galor, P. J. Park, C. A. Walsh, Contrasting somatic mutation patterns in aging human neurons and oligodendrocytes. Cell 187, 1955–1970.e23 (2024).

22. A. Y. Huang, E. A. Lee, Identification of Somatic Mutations From Bulk and Single-Cell Sequencing Data. Front Aging 2, 800380 (2021).

23. B. Milholland, X. Dong, L. Zhang, X. Hao, Y. Suh, J. Vijg, Differences between germline and somatic mutation rates in humans and mice. Nat Commun 8, 15183 (2017).

24. N. Spisak, M. de Manuel, W. Milligan, G. Sella, M. Przeworski, The clock-like accumulation of germline and somatic mutations can arise from the interplay of DNA damage and repair. PLoS Biol 22, e3002678 (2024).

25. H. E. Machado, E. Mitchell, N. F. Øbro, K. Kübler, M. Davies, D. Leongamornlert, A. Cull, F. Maura, M. A. Sanders, A. T. J. Cagan, C. McDonald, M. Belmonte, M. S. Shepherd, F. A. Vieira Braga, R. J. Osborne, K. Mahbubani, I. Martincorena, E. Laurenti, A. R. Green, G. Getz, P. Polak, K. Saeb-Parsy, D. J. Hodson, D. G. Kent, P. J. Campbell, Diverse mutational landscapes in human lymphocytes. Nature 608, 724–732 (2022).

26. C. C. Billingsley, D. E. Cohn, D. G. Mutch, J. A. Stephens, A. A. Suarez, P. J. Goodfellow, Polymerase ɛ (POLE) mutations in endometrial cancer: clinical outcomes and implications for Lynch syndrome testing. Cancer 121, 386–394 (2015).

27. A. McKenna, G. M. Findlay, J. A. Gagnon, M. S. Horwitz, A. F. Schier, J. Shendure, Whole-organism lineage tracing by combinatorial and cumulative genome editing. Science 353, aaf7907 (2016).

28. D. Yang, M. G. Jones, S. Naranjo, W. M. Rideout 3rd, K. H. J. Min, R. Ho, W. Wu, J. M. Replogle, J. L. Page, J. J. Quinn, F. Horns, X. Qiu, M. Z. Chen, W. A. Freed-Pastor, C. S. McGinnis, D. M. Patterson, Z. J. Gartner, E. D. Chow, T. G. Bivona, M. M. Chan, N. Yosef, T. Jacks, J. S. Weissman, Lineage tracing reveals the phylodynamics, plasticity, and paths of tumor evolution. Cell 185, 1905–1923.e25 (2022).

29. J. Choi, W. Chen, A. Minkina, F. M. Chardon, C. C. Suiter, S. G. Regalado, S. Domcke, N. Hamazaki, C. Lee, B. Martin, R. M. Daza, J. Shendure, A time-resolved, multi-symbol molecular recorder via sequential genome editing. Nature 608, 98–107 (2022).

30. P. M. Burgers, Eukaryotic DNA polymerases in DNA replication and DNA repair. Chromosoma 107, 218–227 (1998).

31. 31. Cancer Genome Atlas Network, Comprehensive molecular characterization of human colon and rectal cancer. Nature 487, 330–337 (2012).

32. D. N. Church, S. E. W. Briggs, C. Palles, E. Domingo, S. J. Kearsey, J. M. Grimes, M. Gorman, L. Martin, K. M. Howarth, S. V. Hodgson, NSECG Collaborators, K. Kaur, J. Taylor, I. P. M. Tomlinson, DNA polymerase ε and δ exonuclease domain mutations in endometrial cancer. Hum Mol Genet 22, 2820–2828 (2013).

33. 33. Cancer Genome Atlas Research Network, C. Kandoth, N. Schultz, A. D. Cherniack, R. Akbani, Y. Liu, H. Shen, A. G. Robertson, I. Pashtan, R. Shen, C. C. Benz, C. Yau, P. W. Laird, L. Ding, W. Zhang, G. B. Mills, R. Kucherlapati, E. R. Mardis, D. A. Levine, Integrated genomic characterization of endometrial carcinoma. Nature 497, 67–73 (2013).

34. E. Heitzer, I. Tomlinson, Replicative DNA polymerase mutations in cancer. Curr Opin Genet Dev 24, 107–113 (2014).

35. L. Bar-Peled, N. Kory, Principles and functions of metabolic compartmentalization. Nat Metab 4, 1232–1244 (2022).

36. J. D. Watson, F. H. Crick, Genetical implications of the structure of deoxyribonucleic acid. Nature 171, 964–967 (1953).

37. F. Jacob, Evolution and tinkering. Science 196, 1161–1166 (1977).

38. N. Takahata, Molecular clock: an anti-neo-Darwinian legacy. Genetics 176, 1–6 (2007).

39. R. E. Handsaker, S. Kashin, N. M. Reed, S. Tan, W.-S. Lee, T. M. McDonald, K. Morris, N. Kamitaki, C. D. Mullally, N. R. Morakabati, M. Goldman, G. Lind, R. Kohli, E. Lawton, M. Hogan, K. Ichihara, S. Berretta, S. A. McCarroll, Long somatic DNA-repeat expansion drives neurodegeneration in Huntington’s disease. Cell 188, 623–639.e19 (2025).

40. G. Almogy, M. Pratt, F. Oberstrass, L. Lee, D. Mazur, N. Beckett, O. Barad, I. Soifer, E. Perelman, Y. Etzioni, M. Sosa, A. Jung, T. Clark, E. Trepagnier, G. Lithwick-Yanai, S. Pollock, G. Hornung, M. Levy, M. Coole, T. Howd, M. Shand, Y. Farjoun, J. Emery, G. Hall, S. Lee, T. Sato, R. Magner, S. Low, A. Bernier, B. Gandi, J. Stohlman, C. Nolet, S. Donovan, B. Blumenstiel, M. Cipicchio, S. Dodge, E. Banks, N. Lennon, S. Gabriel, D. Lipson, Cost-efficient whole genome-sequencing using novel mostly natural sequencing-by-synthesis chemistry and open fluidics platform, bioRxiv (2022). 10.1101/2022.05.29.493900.

41. M. Kokoris, R. McRuer, M. Nabavi, A. Jacobs, M. Prindle, C. Cech, K. Berg, T. Lehmann, C. Machacek, J. Tabone, J. Chandrasekar, L. McGee, M. Lopez, T. Reid, C. Williams, S. Barrett, A. Lehmann, M. Kovarik, R. Busam, S. Miller, B. Banasik, B. Kesic, A. Arryman, M. Rogers-Peckham, A. Kimura, M. LeProwse, M. Wolfin, S. Kritzer, J. Leadbetter, M. Babazadeh, J. Chase, G. Thiessen, W. Lint, D. Goodman, D. O’Connell, N. Lumanpauw, J. Hoffman, S. Vellucci, K. Collins, J. Vellucci, A. Taylor, M. Murphy, M. Lee, M. Corning, Sequencing by Expansion (SBX) - a novel, high-throughput single-molecule sequencing technology, bioRxiv (2025). 10.1101/2025.02.19.639056.

42. Y. Arimura, R. M. Shih, R. Froom, H. Funabiki, Structural features of nucleosomes in interphase and metaphase chromosomes. Mol Cell 81, 4377–4397.e12 (2021).

43. C. P. Lapointe, R. Grosely, A. G. Johnson, J. Wang, I. S. Fernández, J. D. Puglisi, Dynamic competition between SARS-CoV-2 NSP1 and mRNA on the human ribosome inhibits translation initiation. Proc Natl Acad Sci U S A 118 (2021).

44. J. Dauparas, I. Anishchenko, N. Bennett, H. Bai, R. J. Ragotte, L. F. Milles, B. I. M. Wicky, A. Courbet, R. J. de Haas, N. Bethel, P. J. Y. Leung, T. F. Huddy, S. Pellock, D. Tischer, F. Chan, B. Koepnick, H. Nguyen, A. Kang, B. Sankaran, A. K. Bera, N. P. King, D. Baker, Robust deep learning-based protein sequence design using ProteinMPNN. Science 378, 49–56 (2022).

45. J. Jumper, R. Evans, A. Pritzel, T. Green, M. Figurnov, O. Ronneberger, K. Tunyasuvunakool, R. Bates, A. Žídek, A. Potapenko, A. Bridgland, C. Meyer, S. A. A. Kohl, A. J. Ballard, A. Cowie, B. Romera-Paredes, S. Nikolov, R. Jain, J. Adler, T. Back, S. Petersen, D. Reiman, E. Clancy, M. Zielinski, M. Steinegger, M. Pacholska, T. Berghammer, S. Bodenstein, D. Silver, O. Vinyals, A. W. Senior, K. Kavukcuoglu, P. Kohli, D. Hassabis, Highly accurate protein structure prediction with AlphaFold. Nature 596, 583–589 (2021).

46. R. Poplin, V. Ruano-Rubio, M. A. DePristo, T. J. Fennell, M. O. Carneiro, G. A. Van der Auwera, D. E. Kling, L. D. Gauthier, A. Levy-Moonshine, D. Roazen, K. Shakir, J. Thibault, S. Chandran, C. Whelan, M. Lek, S. Gabriel, M. J. Daly, B. Neale, D. G. MacArthur, E. Banks, Scaling accurate genetic variant discovery to tens of thousands of samples, bioRxiv (2017). 10.1101/201178.

47. H. Li, R. Durbin, Fast and accurate short read alignment with Burrows-Wheeler transform. Bioinformatics 25, 1754–1760 (2009).

48. H. Li, A statistical framework for SNP calling, mutation discovery, association mapping and population genetical parameter estimation from sequencing data. Bioinformatics 27, 2987–2993 (2011).

49. W. S. DeWitt, L. Zhu, M. R. Vollger, M. E. Goldberg, A. Talenti, A. C. Beichman, K. Harris, Mutyper: Assigning and summarizing mutation types for analyzing germline mutation spectra. J. Open Source Softw. 8, 5227 (2023).

50. I. Virshup, S. Rybakov, F. J. Theis, P. Angerer, F. A. Wolf, anndata: Access and store annotated data matrices. J. Open Source Softw. 9, 4371 (2024).

